# Egocentric vector coding of geometric vertex in the retrosplenial cortex

**DOI:** 10.1101/2023.09.03.556136

**Authors:** Kyerl Park, Yoonsoo Yeo, Kisung Shin, Jeehyun Kwag

## Abstract

Neural representation of the environmental features in a self-centered, egocentric manner is important in constructing an egocentric cognitive map that is critical for goal-directed navigation and episodic memory formation^1^. To create a geometrically detailed egocentric cognitive map, neural representations of edges and vertices of environmental features are needed. While egocentric neural representations of edges, like egocentric boundary vector cells^2–6^ and border cells exist^7^, those of vertices are unknown. Here we report that single neurons in the granular retrosplenial cortex (RSC) generate spatial receptive fields exclusively near the vertices of environmental geometries during free exploration, which we termed vertex cells. Each spatial receptive field of vertex cells occurred at a specific orientation and distance relative to the animal, tuned by head direction, indicating an egocentric vector coding of the vertex. The removal of physical boundaries that define the environmental geometry abolished egocentric vector coding vertex cells. Moreover, goal-directed navigation selectively strengthened the egocentric vertex vector coding at the vertex near the goal location. Overall, our results suggest that egocentric vector coding of vertex by granular RSC neurons help to construct a geometrically detailed egocentric cognitive map that guides goal-directed navigation.

## Main Text

Cognitive map theory proposes that the brain represents spatial information for efficient spatial navigation in the complex environment^8^. Two distinct types of spatial reference frames are believed to be used in spatial navigation: Allocentric and egocentric spatial reference frames^1^. While allocentric neural representations of an animal’s location, orientation, or distance relative to environmental features are well characterized^9–16^, egocentric neural representations of environmental features relative to the animal’s current location and head direction have only recently been discovered^1–7,17,18^. Interestingly, many of the egocentric spatial-selective neurons use vector coding of space^1^ where neurons spike to generate spatial receptive fields when spatial structures are at a certain distance and orientation relative to the animal, tuned by its head direction^1^. The egocentric vector coding cells include boundary vector cells^2–6^, border cells^7^, center bearing/distance cells^17^, and object vector cells^4^. These neurons may collectively contribute to constructing an egocentric cognitive map of the spatial layout of the external world that is important in guiding goal-directed navigation^19^ and episodic memory^1^.

The environmental features in the external world are highly geometric in Euclidean space, which are represented by combinations of edges and vertices. Therefore, an egocentric neural representation of edges and vertices should help the efficient construction of a geometrically detailed egocentric cognitive map of egocentric space that can guide movement transition^20^ for goal-directed navigation^19^. Although egocentric vector coding of edges has been reported in the form of boundary vector or border cells^2–7^, egocentric neural representation of vertices has not been identified yet.

The retrosplenial cortex (RSC) is a good neural circuit candidate for studying egocentric neural representation of vertices. RSC is well established to serve critical roles in goal-directed navigation^19,21–26^, egocentric spatial navigation^2,7,27,28^, episodic memory, as well as transformation between allocentric and egocentric spatial reference frames^7,27^, all of which require an egocentric cognitive map. Anatomically, the granular RSC has dense reciprocal connections with multiple neural circuits that map allocentric space via spatially-tuned cells, including the hippocampus^27,29,30^ (place cells^9^), medial entorhinal cortex^27,31^ (grid cells^10^), anterior thalamic nuclei^27,29,30^ (head direction cells^12^), and subiculum^27,29,30^ (boundary vector cells^13^). Furthermore, egocentric boundary vector/border cells have already been discovered in the RSC^2,7^.

### Neural representation of geometric vertex in the granular RSC

To investigate whether the neural representation of geometric vertices exists in the RSC, we injected AAV-CaMKII-GCaMP6s into the granular layers of RSC and recorded Ca^2+^ signals from a total of 740 GCaMP6s-expressing excitatory neurons using a miniaturized microscope in nine mice (Fig. 1a, b). From the ΔF/Fo Ca^2+^ signals acquired from the granular RSC neurons, we extracted deconvoluted spikes (Fig. 1b) to plot neural activities while mice freely explored open chambers of different environmental geometries. We used a circular open chamber (radius of 25 cm) and three different shapes of regular polygon open chambers that had distinct number of vertices. The dimensions of regular polygon-shaped open chambers were chosen so that they can be inscribed into the circular open chamber, so as to keep the center-to-vertex distance constant (25 cm) (Fig. 1c). In a circular open chamber, the granular RSC neuronal spikes tessellated along the entire circumference (Fig. 1c, d), which is similar to RSC neuronal responses reported in rats^32^. However, in regular polygon-shaped open chambers, subpopulations of granular RSC neurons spiked exclusively near the vertices (Fig. 1c, d). Even when the regular polygon-shaped open chambers were morphed from square to 24-gon using 8 cm-wide walls while keeping the total perimeter constant across chambers, vertex-selective spiking was still observed (Extended Data Fig. 1). The spatial patterns of vertex-selective RSC neuronal receptive fields (Fig. 1c, d) were different from grid cells, as they had low grid scores^10^ (Extended Data Fig. 2). Also, they were distinctively different from boundary vector cells^13^ or border cells^14^ as subpopulations of RSC neurons fired exclusively near the vertex located at either end of the edge, as shown in the populations responses of RSC neurons along a given edge of the open chambers (Fig. 1e). To quantitatively analyze the vertex-selective neuronal characteristics, we defined vertex score. The vertex score was calculated by taking the normalized mean distance of all spatial bins containing the spatial receptive field to the nearest vertex, which was then weighted by the firing rates in each spatial bin. The resulting vertex scores ranged from 0 to 1, where 0 indicated that the spatial receptive fields were located farthest away from all vertices, and 1 indicated that the spatial receptive fields were located exclusively at any of the vertices. A given RSC neuron was defined as a vertex cell within each chamber if the vertex score exceeded the 99^th^ percentile of the probability distribution of randomly shuffled vertex scores in any of the regular polygon-shaped open chambers (Fig. 1f), and if it showed stable spatial receptive fields during the entire 30-min recording sessions. Based on vertex score analysis, in each chamber, 18%, 19%, and 19% of RSC neurons (130, 140, and 139 out of 740 neurons, respectively) were vertex cells in triangle, square, and hexagon chambers, respectively (Fig. 1f, g), while 4% of RSC neurons (32 out of 740 neurons) sustained vertex cell responses in all three different open chambers (Fig. 1g) (Extended Data Fig. 3). Overall, across the three open chambers, 37% (271 out of 740 neurons) of the imaged RSC neurons were defined as vertex cells (Fig. 1g). Among the imaged granular RSC neuronal populations, a high proportion of granular RSC neurons were vertex cells that spiked exclusively near the vertices while being inhibited at the center of the edge (Fig. 1h). However, subpopulations of non-vertex cells in the granular RSC were found to be border cells (17%, 128 out of 740 neurons, Extended Data Fig. 4) while minor subpopulations of vertex cells showed conjunctive responses to both vertices and borders (6%, 43 out of 740 neurons, Extended Data Fig. 5).

**Fig. 1.**
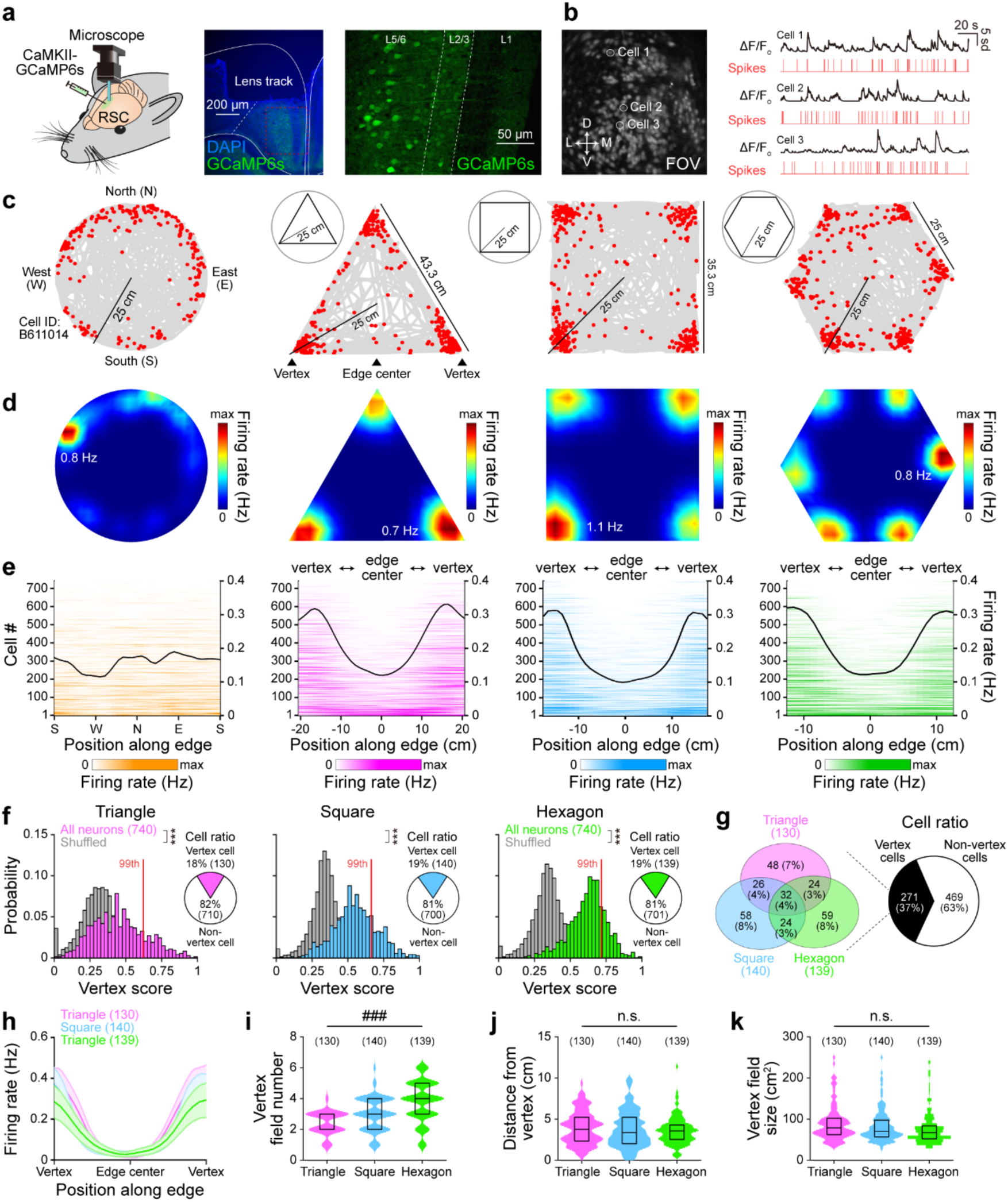
Neural representation of geometric vertices in the retrosplenial cortex. **(a)** Experimental design for Ca^2+^ imaging from mice using miniaturized microscope during free exploration. CaMKII-GCaMP6s injection and a Gradient-index (GRIN) lens implant in the granular retrosplenial cortex (RSC, left). Fluorescent image of DAPI-stained neurons with GRIN lens track (middle) and virally expressed GCaMP6s on the granular RSC neurons (right). (b) Example field of view (FOV) of GCaMP6s-expressing neurons during Ca^2+^ imaging (left). Representative raw Ca^2+^ signals (right, black) from which deconvoluted spikes were extracted (right, red). (**c**, **d**) Representative spike-trajectory plot (c) showing the mouse trajectory (gray line) superimposed with spike locations (red dots) and the corresponding firing rate maps (d) of a single RSC neuron during free exploration in circular, triangle, square, and hexagon open chambers. Firing rate in (d) is color-coded (inset: maximum firing rate). (**e**) RSC neuronal firing rate map along the position on a given edge in each chamber. Black line: mean firing rate of all neurons. (**f**) Probability distributions of vertex scores in triangle (pink), square (blue), and hexagon open chambers (green) plotted together with the randomly shuffled vertex scores for comparison (gray). Red line: 99^th^ percentile of randomly shuffled vertex score distribution. Inset: pie chart showing ratio of vertex cells and non-vertex cells in each open chamber. (**g**) Venn diagram (left) and pie chart (right) and showing vertex cell ratio across three open chambers. (**h**) Average firing rate of vertex cells along the position on a given edge in each chamber. Solid lines: mean firing rate of all vertex cells. Shade: SEM. (**i-k**) Violin plot showing the number of vertex fields (i), distance of vertex fields from vertex (j), and the vertex field size (k) of vertex cells in each open chamber. Box plot shows 25^th^, 50^th^, and 75^th^ percentile values. ***: *p* < 0.001 for unpaired Student’s *t* test (f). ###: *p* < 0.001, n.s.: *p* > 0.05 for one-way ANOVA with *post hoc* Tukey’s test (i-k).

Analyses of spatial receptive fields of vertex cells, termed vertex field henceforth, revealed that they significantly increased in number as the number of environmental vertices within the chamber increased (Fig. 1i). Vertex field locations were ∼ 3 cm proximal to the vertex (Fig. 1j) and vertex field sizes were similarly small (∼ 70 cm^2^) across the chambers (Fig. 1k). Thus, vertex fields covered a small fraction of the edge exclusively near the vertex, which contrasts with border cells (Extended Data Fig. 4). Vertex cell responses were independent of whether the vertex angle was convex (< 180°) or concave (> 180°) (Extended Data Fig. 6). Also, vertex cells’ spatial receptive field properties were robust over time (Extended Data Fig. 7), in different sizes of open chambers (Extended Data Fig. 8), in dark condition (Extended Data Fig. 9), and with visual cue rotation (Extended Data Fig. 10). In addition to deconvoluted spikes, other features of Ca^2+^ data (Ca^2+^ events and raw Ca^2+^ signals) could also capture vertex cell responses (Extended Data Fig. 11). Together, these results show that neurons in the granular RSC provide a robust neural representation of geometric vertices that remains stable over time and across different geometric environments.

### Egocentric vector coding of vertex by granular RSC neurons

Next, to examine the spatial reference frame of vertex cells, we analyzed the tuning of vertex cell’s spikes in either allocentric head direction (Fig. 2a, b) or egocentric bearing relative to the vertex (Fig. 2a, c). Compared with head direction cells in the granular RSC that showed uni-directional allocentric head direction tuning (Extended Data Fig. 12), a given vertex cell’s spikes showed discretization of allocentric head direction tuning at each vertex (Fig. 2d), suggesting a non-allocentric spatial reference frame. When we calculated the egocentric bearing of vertex cell’s spikes as the angle of the mouse head direction relative to the closest vertex, the tuning curves of egocentric bearing were similar across all vertices (Fig. 2d). Thus, vertex cells were tuned to specific egocentric bearing relative to the vertex, indicating that they use an egocentric spatial reference frame. The egocentric reference frame of vertex cells was further characterized by determining the egocentric vertex vector using the egocentric bearing and egocentric distance of spikes^2,3,7^ relative to the closest vertex (Fig. 2e). This egocentric vertex vector was then used to plot an egocentric vertex rate map (Fig. 2f), from which the vector length (VL) of the egocentric vertex vector was analyzed, which quantifies the strength of egocentric vector coding of vertex cells. Using these analyses, a given vertex cell was classified as an egocentric vertex vector cell within each open chamber if the VLs exceeded the 99^th^ percentile of the probability distribution of randomly shuffled VLs (Fig. 2g) (Extended Data Fig. 13). In triangle, square, and hexagon open chambers, 73% (95 out of 130 vertex cells), 74% (104 out of 140 vertex cells), and 76% of vertex cells (105 out of 139 vertex cells) were defined as egocentric vertex vector cells, respectively (Fig. 2g). The VLs of egocentric vertex vector cells were statistically similar across all three open chambers (Fig. 2h) while they were significantly higher than those of non-egocentric vertex vector cells (Fig. 2h). Moreover, their preferred egocentric bearing and egocentric distance sustained across all three open chambers (Fig. 2i, j), over time, in different sizes of chambers, in dark condition, and with visual cue rotation (Extended Data Fig. 14). As mice behavior can be restricted by boundaries and as subpopulations of RSC neurons exhibit direction-selective tuning^28^, whether egocentric vertex vector cell property arise by self-motion was investigated through self-motion referenced rate map analysis^33^. While a small percentage of egocentric vertex vector cells displayed self-motion tuning, the majority exhibited self-motion independence (Extended Data Fig. 15). Thus, our results indicate that subpopulations of neurons in the granular RSC perform robust and stable egocentric vector coding of vertices that is independent of self-motion.

**Fig. 2.**
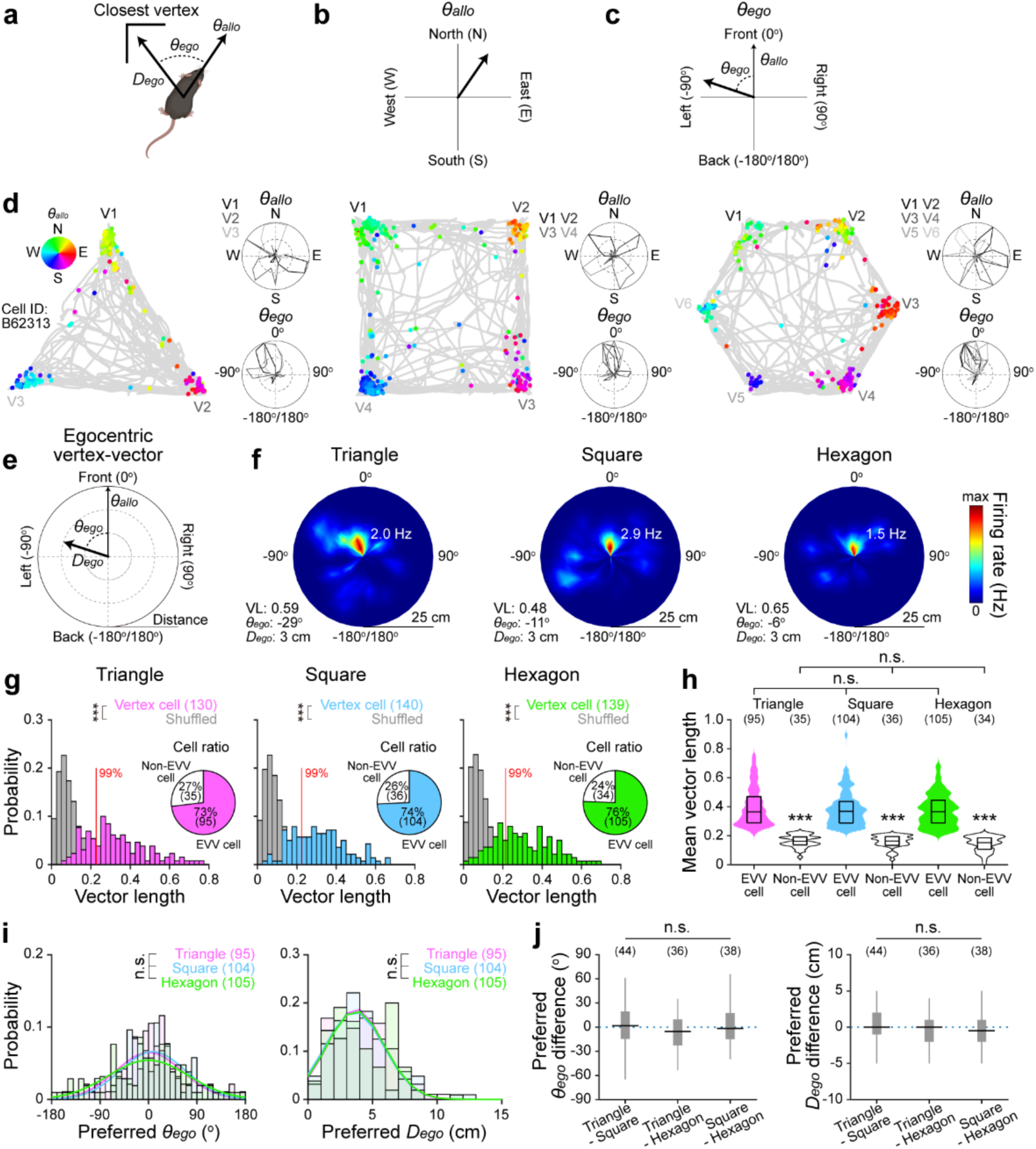
Egocentric vector coding of geometric vertices in the retrosplenial cortex. (**a-c**). Illustration of the mouse allocentric head direction (*θallo*), egocentric bearing (*θego*), and egocentric distance (*Dego*) relative to the closest vertex from the current location (a). *θallo* (b) and *θego* (c, 0° when the mouse faces a vertex). (**d**) Representative spike-trajectory plot showing the mouse trajectory (gray line) superimposed with spike locations (colored dots) of a single vertex cell in the granular retrosplenial cortex during free exploration in triangle, square, and hexagon open chambers. Each spike is color-coded according to the corresponding *θallo* (inset). Allocentric head direction tuning curves and egocentric bearing tuning curves are color-coded for the corresponding closest vertex (Vi: *i*th vertex) in all open chambers. (**e**) Illustration of an egocentric vertex vector plot. (**f**) The corresponding egocentric vertex rate map of the same vertex cell in (d). Firing rate is color-coded. Insets: mean resultant vector length (VL), preferred *θego*, preferred *Dego*, and maximum firing rate. (**g**) Probability distributions of VL in triangle (pink), square (blue), and hexagon open chambers (green) plotted together with the randomly shuffled VLs for comparison (gray). Red line: 99^th^ percentile of randomly shuffled VLs distribution. Inset: pie chart showing ratio of egocentric vertex vector cells (EVV cells) and non-EVV cells in each open chamber. (**h**) Violin plot showing mean VLs of EVV cells and non-EVV cells in each open chamber. (**i**) Probability distribution of preferred *θego* (left) and *Dego* (right) of EVV cells in each open chamber. (**j**) Box plot showing differences in preferred *θego* (left) and *Dego* (right) of EVV cells across chambers. Box plot shows 25^th^, 50^th^, and 75^th^ percentile values. \*\*\**p* < 0.001, n.s.: *p* > 0.05 for unpaired (g, h) and paired (j, right) Student’s *t* test. n.s.: *p* > 0.05 for one-way ANOVA with *post hoc* Tukey’s test (h, i, right). n.s.: *p* > 0.05 for Wilcoxon signed-rank test (i, left, j, left).

### Egocentric vector coding of vertex requires physical boundaries

As vertices are formed at the intersections of boundaries that define the geometry of an environment, the necessity of environmental boundaries in the emergence of egocentric vertex vector cells was investigated by removing the physical boundaries (Fig. 3a). Both vertex-selective firing (Fig. 3b, c) and egocentric vector coding of vertices (Fig. 3d-f) were significantly weakened in the absence of physical boundaries. However, when the physical boundaries were re-introduced, egocentric vector coding of vertices was fully restored (Fig. 3a-f), as shown by the increase in firing rates at vertices (Fig. 3b), vertex scores (Fig. 3c), VLs in egocentric vertex rate map (Fig. 3a, d), and preferred egocentric distance and egocentric bearing (Fig. 3e, f), underscoring the requirement of physical boundaries in adaptive egocentric vertex vector coding of environmental geometry. A simple *in silico* computational model of RSC neuron receiving subthreshold synaptic inputs carrying boundary information could simulate vertex-selective firing characteristics (Extended Data Fig. 16). Thus, integration of spatial information conveying environmental boundaries could be one of the possible input sources in the emergence of vertex cell responses.

**Fig. 3.**
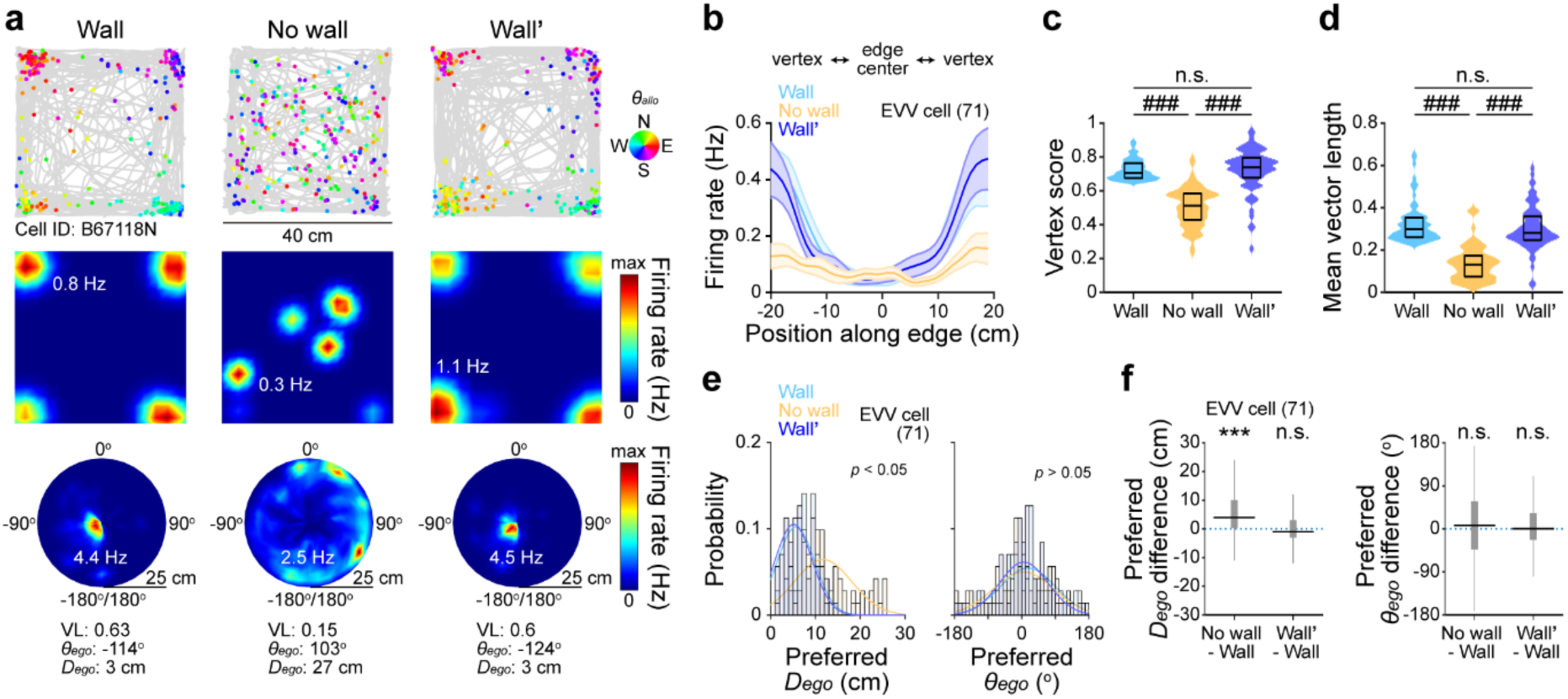
Requirement of environmental physical boundaries for egocentric vector coding of vertices in the retrosplenial cortex. (**a**) Representative spike-trajectory plot (top) showing the mouse trajectory (gray line) superimposed with spike locations (colored dots) and the corresponding firing rate maps (middle) of a single egocentric vertex vector (EVV) cell in the granular retrosplenial cortex (RSC) during free exploration in a square open chamber with (Wall), without (No wall), and with re-introduction of physical boundaries (Wall’). Each spike is color-coded according to the corresponding allocentric head direction (*θallo*) (a, inset, North: N, East: E, South: S, and West: W). Firing rate is color-coded (inset: maximum firing rate). The corresponding egocentric vertex rate map (bottom) of the same EVV cell. Firing rate is color-coded. Insets: mean resultant vector length (VL), preferred egocentric bearing (*θego*), preferred egocentric distance (*Dego*), and maximum firing rate. (**b**) Average firing rate of EVV cells along the position on a given edge in each chamber (Wall: blue, No wall: orange, and Wall’: dark blue). Solid lines: mean firing rate of all EVV cells. Shade: SEM. (**c**) Violin plot showing vertex scores of EVV cells in each chamber. (**d**) Violin plot showing mean VLs of EVV cells in each open chamber. (**e**) Probability distribution of preferred *Dego* (left) and *θego* (right) of EVV cells in each open chamber. (**f**) Box plot showing differences in preferred *Dego* (left) and *θego* (right) and of EVV cells across chambers. Box plot shows 25^th^, 50^th^, and 75^th^ percentile values. ###: *p* < 0.001, n.s.: *p* > 0.05 for One-way ANOVA with *post hoc* Tukey’s test (c, d, e, left). \*\*\**p* < 0.001, n.s.: *p* > 0.05 for Wilcoxon signed-rank test (e, right, f, right). \*\*\**p* < 0.001, n.s.: *p* > 0.05 for paired Student’s *t* test (f, left).

### Goal-directed navigation strengthens egocentric vector coding of vertex at goal location

RSC has been proposed to be important in goal-directed navigation^19,21–26^ using egocentric cognitive map^1^. To directly test egocentric vertex vector cell’s contribution to goal-directed navigation, we designed a goal-directed navigation task in a complex environment consisting of a square open chamber with three connecting inner walls (Fig. 4a), in which mice was trained to learn the goal location (top-left vertex, V1) where a reward was given (Fig. 4a, b). We monitored the egocentric vertex vector cells responses during free exploration in the complex environment before and after the goal location learning. Before goal-directed navigation, egocentric vertex vector cells responded to both the vertices of the square open chamber and the inner walls (Fig. 4c) with the same preferred egocentric bearing and distance across the vertices (Extended Data Fig. 17). However, after learning the goal location, spike firing rate (Fig. 4d, e) and VLs of the same egocentric vertex vector cells significantly increased compared to before learning exclusively at V1 (Fig. 4d, f, g). Moreover, after goal location learning, egocentric vertex vector cell’s preferred egocentric distance at V1 significantly increased while egocentric bearings were sustained (Fig. 4h). The goal location learning-induced changes in egocentric vector coding of the vertex was independent of the mice occupancy time at the goal location (Extended Data Fig. 18). Overall, these results indicate that goal location learning induces strengthening of egocentric vector coding of vertices in the complex environment to guide efficient goal-directed navigation.

**Fig. 4.**
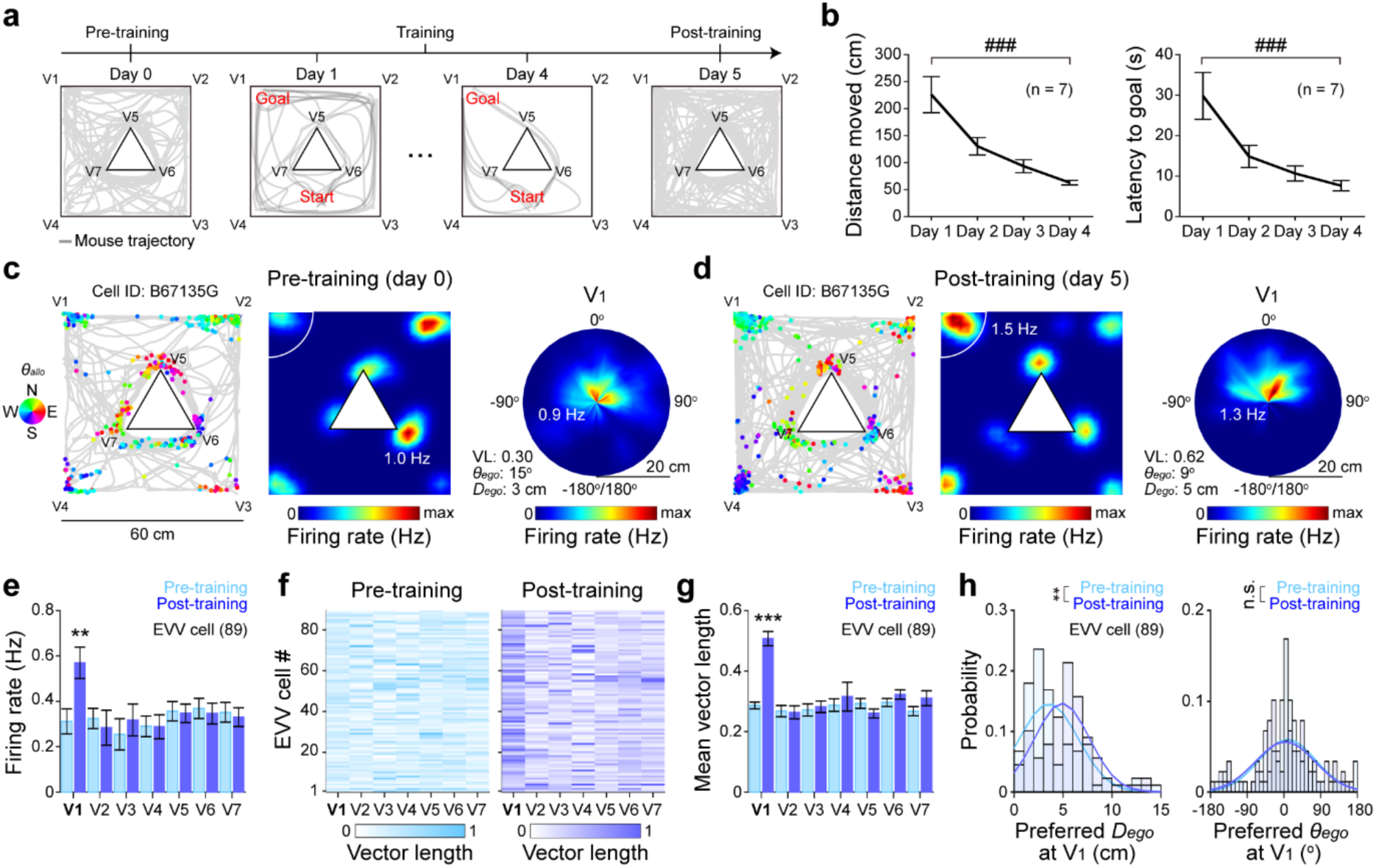
Strengthening of egocentric vector coding of geometric vertices in the retrosplenial cortex by goal-directed navigation. (**a**) Experimental timeline for goal-directed navigation task (top). Representative mouse trajectories (gray line) during the goal-directed navigation task in a square open chamber with a triangle-shaped inner walls (Vi: *i*th vertex) on pre-training, training, and post-training sessions (bottom). (**b**) Averaged trajectory distance (left) and latency (right) from start position to goal location during training sessions (n = 7 mice). (**c**, **d**) Representative spike-trajectory plot (left) showing the mouse trajectory (gray line) superimposed with spike locations (colored dots) and the corresponding firing rate maps (middle) of a single egocentric vertex vector (EVV) cell in the granular retrosplenial cortex during free exploration in the environment before (c, Pre-training) and after (d, Post-training) training sessions. Each spike is color-coded according to the corresponding allocentric head direction (*θallo*) (inset, North: N, East: E, South: S, and West: W). Firing rate is color-coded (inset: maximum firing rate). The corresponding egocentric vertex rate map (right) of the same EVV cell at the vertex near goal location (V1, white line in firing rate map). Firing rate is color-coded. Insets: mean resultant vector length (VL), preferred egocentric bearing (*θego*), preferred egocentric distance (*Dego*), and maximum firing rate. (**e-g**) Average firing rate (e), VL map (f), and mean VL (g) of all EVV cells at each vertex in each condition (Pre-training: blue, Post-training: dark blue). VL in (g) is color-coded. (**h**) Probability distribution of preferred *Dego* (left) and *θego* (right) of EVV cells at V1 in each condition. Bar graph shows mean ± SEM. ###: *p* < 0.001 for One-way ANOVA with *post hoc* Tukey’s test (b). \*\*\**p* < 0.001, \*\**p* < 0.01 for paired Student’s *t* test (e, g, h, left). n.s.: *p* > 0.05 for Wilcoxon signed-rank test (h, right).

## Discussion

In our study, we present a novel type of spatially-tuned neuron in the granular RSC that represents geometric vertices of the environmental features in an egocentric manner (Fig. 1, 2). Egocentric vertex vector cells required physical boundaries (Fig. 3) and goal location learning through goal-directed navigation selectively strengthened the egocentric vector coding of the vertex at the goal location (Fig. 4). Together, these results suggest that egocentric vertex vector cell identified in our study may help construct a geometrically detailed egocentric cognitive map that can guide goal-directed navigation.

A single egocentric vertex vector cell had distinctly small spatial receptive field sizes (∼ 70 cm^2^) (Fig. 1) compared to other spatial selective neurons ^10,14,16^ and generated multiple spatial receptive fields (Fig. 1). Additionally, the sizes and locations of egocentric vertex vector cell’s spatial receptive fields were robust across different environmental geometries (Fig. 1, 2), unlike other spatial navigation neurons that exhibit spatial receptive field remapping^20,34,35^. Thus, egocentric vertex vector cells are ideal for providing a stable and robust one-to-one egocentric mapping of geometric vertices present in the environmental features (Fig. 1, 2).

In the absence of physical boundaries, egocentric vertex vector cells lost their sensitivity to vertex as well as egocentric tuning (Fig. 3), supporting the requirement of physical boundaries in egocentric vertex vector cell generations. Indeed, our simple *in silico* model demonstrated that subthreshold synaptic inputs carrying boundary vector information^27^ are sufficient to simulate vertex cell responses (Extended Data Fig. 16). However, granular RSC neurons receive direct projections from multiple neural circuits processing spatial navigation^27,29–31^ as well as from the visual cortex^19^. Also, granular RSC neurons may receive inputs from multi-directional head direction cells^36^ and egocentric boundary vector/border cell^2,7^ in the dysgranular RSC. Thus, considerations of these inputs in future studies may help elucidate how egocentric vertex vector cell responses arise and how they relate to known functions of RSC including goal-directed navigation^19,21–26^, visual cues/landmarks processing^36–38^, episodic spatial memory^27^, and motor planning^19,27^.

Egocentric vector coding of vertices was strengthened selectively at goal location after goal-directed navigation (Fig. 4), which is in line with RSC’s role in goal-directed navigation^19,21–26^. Thus, egocentric vertex vector cells might contribute to goal-directed navigation by providing egocentric vectorial information of the goal location to hippocampal place cells or neurons in the lateral entorhinal cortex, so as to help them construct the goal-oriented vectorial map^4,39–41^. Also, the strengthening of egocentric vector coding after goal location learning may contribute to goal-related reorganization of allocentric^42–48^ and egocentric cognitive maps^4^. Considering that granular RSC output is well known to project to motor cortex^19^, egocentric vertex vector cells may also help fine-tune motor output that guides movements toward the goal location, which requires further investigation.

Despite these results, there are many limitations to our study. Firstly, we based our results on Ca^2+^ imaging data, which has lower temporal resolution than spike recordings. Moreover, some corner-selective neurons tuned by turning direction^20^ and internal states^49^ have been reported previously. Therefore, careful dissociation of self-motion and internal states in vertex cell analysis with *in vivo*-recorded spikes may be needed in the future. Nevertheless, egocentric vertex vector cells identified here provide insight into the egocentric neural coding of geometric features in the external environment. These neurons could serve pivotal functions in constructing a geometrically detailed and spatial memory-based egocentric cognitive map that is updated continually in changing environment, which in turn could guide adaptive goal-directed navigation and episodic spatial memory in the complex environment.

## Acknowledgments

We thank Dr. Michael M. Kohl (University of Glasgow, UK) for advice on setting up *in vivo* Ca^2+^ imaging.

## Funding

National Research Foundation of Korea grant NRF-2019M3E5D2A01058328 (J.K.)

## Author contributions

Conceptualization: J.K.

Methodology: J.K., K.P., Y.Y., K.S.

Investigation: J.K., K.P., Y.Y., K.S.

Visualization: J.K., K.P., Y.Y., K.S.

Funding acquisition: J.K.

Project administration: J.K.

Supervision: J.K.

Writing – original draft: J.K., K.P.

Writing – review & editing: J.K., K.P., Y.Y., K.S.

## Competing interests

“Authors declare that they have no competing interests.”

## Data and materials availability

“All data are available in the main text or the supplementary materials.”

## Supplementary Materials for Egocentric vector coding of geometric vertex in the retrosplenial cortex

### Materials and Methods

#### Animals

A total of nine C57BL/6 mice (3 – 6 months old; DBL, South Korea) were used for *in vivo* Ca^2+^ imaging during free exploration with the approval of Institutional Animal Care and Use Committee (IACUC) at Korea University (KUIACUC-2020-0099, KUIACUC-2021-0052). All mice were individually housed in a temperature- and humidity-controlled vivarium, which was kept in a 12 h/12 h light/dark cycle. All experiments were performed during the dark cycle. Food and water were available *ad libitum* unless it is mentioned otherwise.

#### Virus and stereotaxic surgery

For surgical procedure, mice were deeply anesthetized using 2% isoflurane (2 ml/min flow rate) and head-fixed into a stereotaxic frame (51730D, Stoelting Co., USA). AAV9-CaMKII-GCaMP6s^1^ (#107790, Addgene, USA) in solution (500 nl per injection site, 2.5 × 10^13^ virus molecules/ml diluted with saline at 3:1 ratio) was injected into three sites along the dorsoventral axis of the retrosplenial cortex (RSC) (AP: –2.5 mm, ML: –0.3 mm, DV: –0.8, –0.6, and –0.4 mm, Fig. 1a) using a Hamilton syringe (#87930, Hamilton, USA) controlled by an automated stereotaxic injector (100 nl/min; #53311, Stoelting Quintessential Injector, Stoelting Co., USA). The syringe was left at the injection coordinates for more than 5 min to allow viral diffusion.

After viral injection, a Gradient-index (GRIN) lens (1 mm diameter, 4 mm length; #1050-004605, Inscopix, USA) was implanted to RSC (AP: –2.5 mm; ML: –0.3 mm; DV: –0.5 mm) and fixed to the skull with dental cement (Self Curing, Vertex, Netherlands). After two weeks of recovery time following the surgery, mice were deeply anesthetized using 2% isoflurane (2 ml/min flow rate) to attach a magnetic baseplate (#1050-004638, Inscopix, USA), on top of which miniaturized microscope (nVoke, Inscopix, USA) was mounted. If GCaMP6s-expressing neurons were visible in the field of view (FOV), baseplate was fixed to the skull using dental adhesive resin cement (Super bond, Sun Medical, Japan) to perform *in vivo* longitudinal Ca^2+^ imaging of the same FOV. After baseplanting, at least one week of recovery time was allowed before performing *in vivo* Ca^2+^ imaging.

#### Behavior and chamber design

Prior to *in vivo* Ca^2+^ imaging during free exploration, mice were adapted to the head-mounted miniaturized microscope connected to the commutator systems (Inscopix, USA) for at least five days. All experiments were performed under the luminescence of 30 Lux unless it is mentioned otherwise.

In Fig. 1–2, each mouse was allowed to freely explore four different shapes of open chambers with different geometry: circular open chamber (radius: 25 cm, height: 30 cm), a regular triangle open chamber (edge: 43.3 cm, height: 30 cm), a square open chamber (edge: 35.3 cm, height: 30 cm), and a regular hexagon open chamber (edge: 25 cm, height: 30 cm). The dimensions of regular polygon-shaped open chambers were designed to be inscribed into a circular open chamber of 25 cm-radius to keep the center-to-vertex distance in each regular polygon-shaped open chamber constant (25 cm). In Fig. 3, the square open chamber (edge: 40 cm, height: 30 cm) was inverted to create an arena with no physical boundaries.

In Fig. 4 and in Extended Data Fig. 17-18, goal-directed navigation task was performed. For this, mice were food-restricted until they reached 80% – 85% of their free-feeding weight^2^. A day before (pre-training session) the goal-directed navigation training started, each mouse was allowed to freely explore a complex environment consisting of a square open chamber (edge: 60 cm, height: 30 cm) with three walls inserted within the chamber that made a regular triangle-shape (edge: 20 cm, height: 20 cm) for 30 min. Goal directed navigation training was performed over four days where each mouse was trained to forage in the chamber which had a food reward (Froot Loops, Kellogg’s, USA) located close to the top-left vertex (V1) of the square chamber (Fig. 4A). Each training began by placing the mouse at the start position and was ended when the mouse reached the goal location and consumed the Froot Loops. Training was repeated 10 times per day. A day after the end of goal-directed navigation training (post-training session), each mouse was allowed to freely explore the same chamber for 30 min in the complex environment without a reward/goal. No Froot Loops were presented during neither pre-training nor post-training sessions.

In Extended Data Fig. 1, to keep the total perimeter of the environmental geometry constant across different shapes of environmental geometry, five different regular polygon-shaped open chambers were configured using 24 identical pieces of rectangular foam board (width: 8 cm, height: 30 cm, total perimeter 192 cm). The shapes of the open chambers were continuously transformed or “morphed” from a square (edge: 48 cm) to hexagon (edge: 32 cm), octagon (edge: 24 cm), 12-gon (edge: 16 cm), and 24-gon open chamber (edge: 8cm) in a series. In Extended Data Fig. 6, a regular concave open chamber (edge: 25 cm, height: 30 cm) with three different vertex angles (60°, 120 °, and 240 °) was used. In Extended Data Fig. 7 and 14, a square open chamber (edge: 35.3 cm, height: 30 cm) was used. In Extended Data Fig. 8 and 14, two different sizes of square open chambers were used: one with 35.3 cm-edge and the other with 60 cm-edge, both with 30 cm-height. In Extended Data Fig. 9 and 14, a square open chamber (edge: 35.3 cm, height: 30 cm) was used in the presence (30 Lux) or absence of visible light (0 Lux). In Extended Data Fig. 10 and 14, a square open chamber (edge: 35.3 cm, height: 30 cm) with circular and striped patterned visual cues (10 cm x 10 cm for each) was used. Visual cues were attached to one side of the chamber first (Extended Data Fig. 10a, middle, 14h), which was then rotated by 90° (Extended Data Fig. 10a, right, 14i).

Each mouse was allowed to freely explore in each open chamber or arena for 30 min, after which each open chamber was cleaned using 70% ethanol to eliminate any odors. Froot Loops were randomly scattered on the floor of the open chambers to facilitate the full coverage of behavior trajectory on the testing arena. Mouse behavior and trajectories were recorded by a video camera placed on the ceiling above the chambers or arenas at a sampling rate of 7.5 Hz. All behavior data including (x, y) coordinates, velocity, and head direction were analyzed using deep learning-based body points detection methods included in Ethovision XT 16 program (Noldus, Netherlands). The x, y coordinates and velocity of the mouse were determined based on the body center point of the mouse. Head direction was measured based on the nose point and the body center point. Epochs of data with velocity slower than 2 cm/s were excluded from data analysis, which ensured analysis of data during active exploration.

#### *In vivo* Ca^2+^ imaging and signal processing

*In vivo* Ca^2+^ imaging data from GCaMP6s-expressig RSC neurons were acquired using nVoke acquisition software (Inscopix, USA) at a sampling rate of 20 Hz (exposure time: 50 ms). An optimal LED power (0.1 – 0.3 mW/mm^2^) and digital focus (0 – 1000) was selected for each mouse based on GCaMP6s expression on neurons in the FOV and the same parameters were used for each mouse for longitudinal imaging.

Ca^2+^ image processing was performed using Inscopix Data Processing Software (Inscopix, USA), following other studies^3,4^. For all data, motion correction was applied using an image registration method^3,5^. Mean fluorescence signal value (F) of each pixel across the entire Ca^2+^ image video was used to calculate the reference, termed Fo. Then, changes in fluorescence signal value (△F) of each pixel were normalized by Fo to acquire △F/Fo. Putative neurons and their Ca^2+^ signals were isolated using automated cell-segmentation algorithm on △F/Fo data, which is based on principal component analysis-independent component analysis^3,6^.

Putative spikes were inferred from △F/Fo signal using spike deconvolution algorithm called “Online Active Set methods to Infer Spikes”^4,7–9^. Spikes with an amplitude less than 3 signals-to-noise ratio of deconvolved Ca^2+^ signals were removed from the data. Ca^2+^ transient events (Extended Data Fig. 11b-d) were defined using Ca^2+^ event detection algorithm which detects the large amplitude peaks (3 standard deviation (sd) from △F/Fo signal) with slow decays (minimum: 0.2 s) by thresholding at the local maxima of the △F/Fo. For △F/Fo-based analysis (Extended Data Fig. 11e, f), sample data with the value of △F/Fo lower than 1.5 sd was excluded.

#### Histology

To confirm the position of GRIN lens track and GCaMP6s expression in RSC neurons after Ca^2+^ imaging, the brain was removed from the mice which were deeply anesthetized using Avertin and perfused with 10 ml chilled 4% paraformaldehyde (158127, Sigma-Aldrich, USA) transcardially. The brain was fixed overnight in 4% paraformaldehyde and transferred to 30% sucrose solution in phosphate-buffered saline (PBS) for cryoprotection for 24 – 48 h. Brains were frozen in Optimum Cutting Temperature (OCT) compounds (Tissue-Tek O.C.T. Compound, SAKURA, Japan) at –50°C prior to cryosectioning in 50 μm-thick coronal slices on a cryostat (YD-2235, Jinhua Yidi Medical Appliance Co., China). Slices were washed three times with PBS to wash out the OCT compounds and mounted on slide glasses with an antifade mounting medium with DAPI (Vectashield, Vector Laboratories, USA). Imaging locations were identified based on the position of GRIN lens track. Expression of GCaMP6s in neurons was verified by detecting the fluorescent signal using a confocal microscope (LSM-700, ZEISS, Germany) and fluorescent microscope (DM2500, Leica, Germany).

#### Spatial receptive field and firing rate map analyses

Ca^2+^ imaging data from Inscopix and behavior data from Ethovision XT 16 were synchronized using Noldus-IO box (Mini USB-IO box, Noldus, Netherlands) system, which were then analyzed with custom-made Matlab (Mathworks, USA) code. To analyze the spatial receptive fields of GCaMP6s-expressing RSC neurons, positional occupancy of each mouse within an open chamber was discretized into 2.5 cm x 2.5 cm spatial bins. Within each spatial bin, the number of spikes evoked was divided by the total time spent within the bin to derive firing rate maps. Only spatial bins with more than 3 spikes and in which mice spent more than 133 ms (temporal resolution of behavior data) were used in spatial receptive field analysis. Firing rate maps were smoothed by applying boxcar average (5 x 5 bins) with a Gaussian smoothing kernel (σ = 1.5). A spatial receptive field was defined by detecting the spatial bins of at least 20 cm^2^ that showed firing rates which were above 30% of the maximum firing rate in the smoothed firing rate map, following the definition of spatial receptive fields in other spatial navigational neurons^10,11^ with modification of minimum spatial receptive field size. To analyze spatial stability of RSC neuronal firing rate map over time (Fig. 1–4, Extended Data Fig. 7) and across different conditions (Extended Data Fig. 9, 10), spatial correlation coefficients (%) between firing rate maps were calculated using “Corr2” function in Matlab toolbox where % is defined as below:

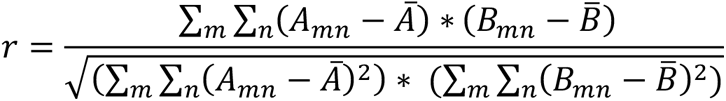

where *m* and *n* are the number of spatial bins for horizontal and vertical axis, respectively, *A*_mn“_ and *B_mn_* are mean firing rates of two firing rate maps for (*m, n*) coordinate spatial bin, respectively, and 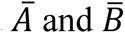 are mean firing rates of two firing rate maps for all spatial bins, respectively. A neuron with % between the first half (0 – 15 min) and the second half (15 – 30 min) of each imaging session greater than 0.5 was included in data analysis following the definition of spatial stability in other spatial navigation neurons^10^.

#### Vertex score analysis

To define and characterize vertex-selective spatial receptive fields, vertex score was introduced. Vertex score was computed by measuring the spatial distribution of a single neuronal spatial receptive fields relative to the vertex, a similar approach used in defining the border scores^10^ of neurons that show border-selective spatial receptive fields. Thus, vertex score (3) of a given neuron was calculated as follows:

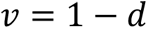

where *d* is the normalized mean distance of all spatial bins, containing spatial receptive fields, to the nearest vertex, which was weighted by the normalized firing rates in each spatial bin. In this way, *v* value ranged between 0 to 1, where *v* = 0 indicated all spatial receptive fields were located farthest away from all vertices and *v* = 1 indicated all spatial receptive fields were located exclusively at any of the vertices. A neuron was defined as vertex cell within each chamber if vertex score for each chamber exceeded the 99^th^ percentile of randomly shuffled vertex score distribution of all imaged RSC neurons. Shuffled vertex score distribution was generated by circularly time-shifting the spike trains of neurons with randomly selected period along the mouse trajectory with 100 repetitions following the method used in previous studies^12,13^. This random time-shuffling preserved the temporal structures of both Ca^2+^ signals and behavior data while disrupted the temporal correlation between them. Random time-shuffling was applied for all randomly shuffled distribution for other metrics.

#### Head direction tuning analysis

To analyze the head direction tuning of a RSC neuronal spikes near each vertex (Fig. 2d, Extended Data Fig. 13a, c, e, inset, top), individual allocentric head direction tuning curves at each vertex were plotted by taking the normalized spike firing rate as a function of allocentric head direction (*θallo*). For each vertex, spikes that were located within the half-length of edge from the corresponding vertex (triangle: 21.65 cm, square: 17.65 cm, and hexagon: 12.50 cm) were used in the tuning curve analysis. In Extended Data Fig. 12, allocentric head direction tuning curves were plotted by taking the normalized firing rate of all spikes in the open chamber as a function of *θallo*. Allocentric head direction tuning curve was plotted using 15° bin and smoothed using a moving average filter.

#### Egocentric vertex vector analysis

To analyze the egocentric bearing of the mouse to vertex (*θego*), the angle of the closest vertex relative to *θallo* at each sample point of behavior data was computed. *θego* was set to 0°, 90°, – 180/180°, and –90°, when the closest vertex is in front, right, back, and left to mice, respectively. Egocentric distance to vertex (*Dego*) was calculated by computing the distance from mice to the closest vertex at each sample point of behavior data. In order to analyze the tuning of egocentric bearing of a RSC neuronal spikes (Fig. 2d, Extended Data Fig. 13a, c, e, inset, bottom), individual egocentric bearing tuning curves at each vertex were plotted by taking the normalized firing rate of spikes as a function of egocentric bearing (*θego*). For each vertex, spikes that were located within the half-length of edge from the corresponding vertex (triangle: 21.65 cm, square: 17.65 cm, and hexagon: 12.50 cm) were used in the tuning curve analysis. Egocentric bearing tuning curve was plotted using 15° bin and smoothed using a moving average filter.

To analyze RSC neuronal spike firing characteristics in terms of *θ_ego_* and *D_ego_*, egocentric vertex rate map was analyzed (Fig. 2, 3, 4, Extended Data Fig. 13, 14, 15, 17, 18). Egocentric vertex rate map was plotted by calculating the firing rate in terms of *θ_ego_* and *D_ego_* in a polar coordinate in polar plot. *θ_ego_* was divided into 15° bins and egocentric distance was divided into 1 cm bins, which yielded 15° x 1 cm bins. Firing rates for each bin was calculated for each neuron, by dividing the number of spikes within a given bin by the whole time spent in the bin. Only the bin in which mice spent more than 133 ms (temporal resolution of behavior data) was included in data analysis. Egocentric vertex rate maps were smoothed using 2D Gaussian kernel σ = 0.7 pixels.

To quantify egocentric vertex vector sensitivity of RSC neurons (Fig. 2, 3, 4, Extended Data Fig. 13, 14, 15, 17, 18), the mean resultant egocentric vertex vector was calculated by adopting the previous definition of egocentric boundary vector^12–14^ with slight modification as:

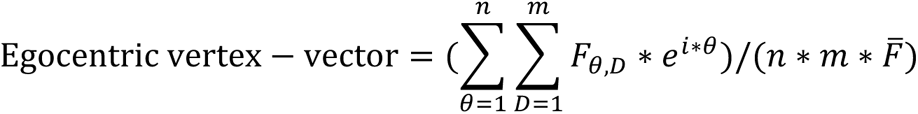

where *θ* indicates the egocentric bearing to vertex, *D* indicates the egocentric distance to vertex, *F_θ,D_*_’_ indicates the firing rate in a given (egocentric bearing ∗ distance) bin, *n* indicates the number of degree bins, *m* indicates the number of distance bins, *i* is the Euler constant, *i* is the imaginary constant, and A 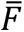 indicates the mean firing rates of all bins.

The vector length of egocentric vertex vector was defined as the absolute value of the mean resultant egocentric vertex vector, which quantifies the strength of egocentric tuning to vertex. Preferred *θego* of egocentric vertex vector was calculated as the mean resultant bearing (MRB):

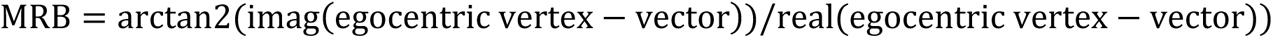

Preferred *Dego* of egocentric vertex vector was estimated by fitting a Weibull distribution^12^ to the firing rates corresponding to the MRB and finding the distance bin with the maximum firing rate.

The vector length of egocentric vertex vector ranged from 0 to 1, where 0 indicated neurons with no egocentric vector coding of vertices and 1 indicated neurons with perfect egocentric vector coding of vertices. A neuron was defined as egocentric vertex vector cell within each chamber if the egocentric vertex vector lengths for each chamber exceeded the 99^th^ percentile of shuffled distribution of the vector length of all imaged neurons.

#### Grid score analysis

To investigate whether spatial receptive fields of RSC neurons have grid cell-like characteristics, grid score was computed (Extended Data Fig. 2) following the same method described in other grid cell studies^15,16^. First, spatial autocorrelograms were calculated as:

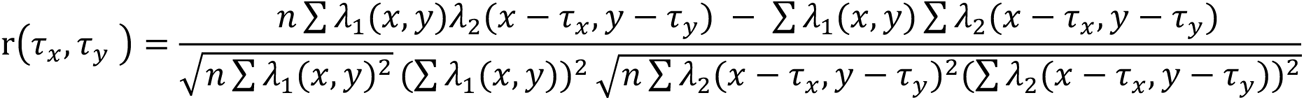

where (*τ_x_,τ_y_*) is the autocorrelation between bins with spatial offset*τ_x_* and *τ_y_*, λ is the firing rate map of a neuron with no smoothing applied, and 2 is the number of overlapping bins in two offset copies of the map. The autocorrelogram was smoothed using boxcar averaging of spatial bins (5 x 5 bins) using a Gaussian smoothing kernel with σ = 1.5. Grid score was calculated by repeatedly rotating the smoothed autocorrelation plot by 6° and the spatial correlation between the rotated autocorrelation plot and the original autocorrelation was calculated as the difference between minimum correlation coefficient for rotations of 60° and 120° and the maximum correlation coefficient for rotations of 30°, 90°, and 150°. A neuron was defined as grid cell within each chamber if grid score for each chamber exceeded the 99^th^ percentile of randomly shuffled grid score distribution of all imaged neurons.

#### Border score analysis

Border cell analysis was performed (Extended Data Fig. 4, 5) following the same method described in the previous study^10^. Putative border fields were defined first by detecting the adjacent spatial receptive field along the border with firing rate higher than 30% of the peak firing rate of the smoothed firing rate map. For all experiments, the maximum value of any single coverage of border field over any environmental border (*C*) was calculated by computing the fraction of spatial bins along the border which was occupied by the spatial receptive fields. The mean firing distance (*e*) indicates the mean distance of all spatial bins, containing receptive fields, to the nearest border, weighted by the normalized firing rate in the spatial bins. e was divided by the maximum distance to borders to obtain e value ranging from 0 to 1. A border score (*b*) for a neuron was defined by comparing *c* and *e*, adopted from the previous research^10^:

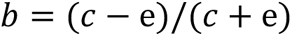

*b* ranged from –1 to 1, where *b =* –1 indicated no spatial receptive fields along any of the borders and *b* = 1 indicated all spatial receptive fields perfectly located along any of borders. A neuron was defined as border cell within each chamber if border score for each chamber exceeded the 99^th^ percentile of randomly shuffled border score distribution of all imaged neurons.

#### Head direction score analysis

To characterize the strength of allocentric head direction tuning of a RSC neuron, head direction score^17^ was computed (Extended Data Fig. 12) based on the mean vector length from circular distributed firing rates in terms of *θallo*. A neuron was defined as head direction cell within each chamber if head direction score for each chamber exceeded the 99^th^ percentile of randomly shuffled head direction score distribution of all imaged neurons.

#### Self-motion referenced rate map analysis

To investigate whether a RSC neuron shows self-motion tuning, self-motion referenced rate maps of the neuron in triangle, square, and hexagon open chambers were analyzed (Extended Data Fig. 15) following the methods used in the previous study^18^. First, for all open chambers, self-motion referenced movement-vectors (Extended Data Fig. 15a) of each mouse at each sample point of behavior data (temporal resolution: 133 ms) were analyzed, which consist of lateral and longitudinal translation of the mouse. Lateral and longitudinal translation of the mouse were computed by converting the velocity and *θallo* of the mouse at each sample point of behavior data to Cartesian coordinates to generate x − and y − displacement values in centimeters per second. Then, self-motion referenced rate map was plotted (Extended Data Fig. 15d, g) by calculating the firing rate in terms of lateral and longitudinal translation of the mouse. Both lateral and longitudinal translations were divided into 1 cm/s x 1 cm/s bins. Firing rates for each bin was calculated for each neuron, by dividing the number of spikes within a given bin by the total time spent in the bin. Only the bins with more than 3 spikes and in which mice spent more than 133 ms (temporal resolution of behavior data) were included in data analysis. Self-motion referenced rate maps were smoothed using 2D Gaussian kernel σ = 0.7 pixels.

To quantify self-motion tuning of RSC neurons, coherence of self-motion referenced rate map of the neuron was calculated by measuring the mean correlation between the firing rate in each bin and the average firing rate of the eight adjacent bins^19^. Coherence of a self-motion referenced rate map ranged from 0 to 1, where 0 indicated neurons with no self-motion tuning and 1 indicated neurons with perfect self-motion tuning. A neuron was defined as self-motion tuned neuron within each chamber if coherence for each chamber exceeded the 99^th^ percentile of shuffled distribution of the coherence of all imaged neurons.

#### *In silico* computational model of vertex cell

To investigate whether RSC vertex cell responses can be simulated simply with a neuron receiving synaptic inputs carrying boundary vector information, *in silico* computational model of RSC and boundary vector cells were constructed (Extended Data Fig. 16). RSC neuron model was adopted from Brennan et al.^20^ (ModelDB accession number 260192), which was a conductance-based simple multi-compartment Hodgkin-Huxley-type neuron model of excitatory neuron in RSC described by the following equation:

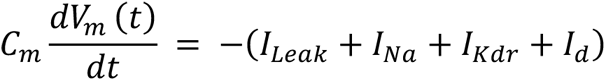

where *C_m_* is the membrane capacitance, *I_leak_* is the leak current, *I_Na_* is the fast sodium channel current, *I_ldr_* is the delayed-rectifier potassium channel current, and *I_d_* is the d-current^21^. The parameters used for the RSC neuron model are shown in Extended Data Table 1, adopted from Brennan et al.^20^.

**Extended Data Table 1.** Membrane properties of in silico RSC neuron model.

| Properties | Values in soma | Values in dendrite |
| --- | --- | --- |
| Capacitance ($\mu\text{F}/\text{cm}^2$) | 1 | 1 |
| Membrane resistance ($k\Omega\text{cm}^2$) | 14.29 | 14.29 |
| Axial resistance ($\Omega\text{cm}$) | 0.000000001 | 200 |
| Maximal conductance of $I_{Leak}$ ($g_{Leak}$, $\text{S}/\text{cm}^2$) | 0.00007 | 0.00007 |
| $g_{Na}$ ($\text{S}/\text{cm}^2$) | 5.5 | 0.81 |
| $g_{Kdr}$ ($\text{S}/\text{cm}^2$) | 0.07 | 0.006 |
| $g_d$ ($\text{S}/\text{cm}^2$) | 0.075 | 0 |

*In silico* computational model of boundary vector cell was built as a conductance-based single-compartment Hodgkin-Huxley-type neuron model where membrane potential of boundary vector cell model was described by the following equation:

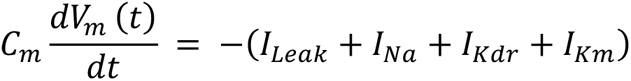

where *C_m_* is the membrane capacitance, *I_leak_* is the leak current, *I_Na_* is the fast sodium channel current, *I_ldr_* is the delayed-rectifier potassium channel current, and *I_d_* is is the M-type potassium channel current^22^. The parameters used for the boundary vector cell model are shown in Extended Data Table 2, adopted from Jang et al.^23^.

**Extended Data Table 2.**
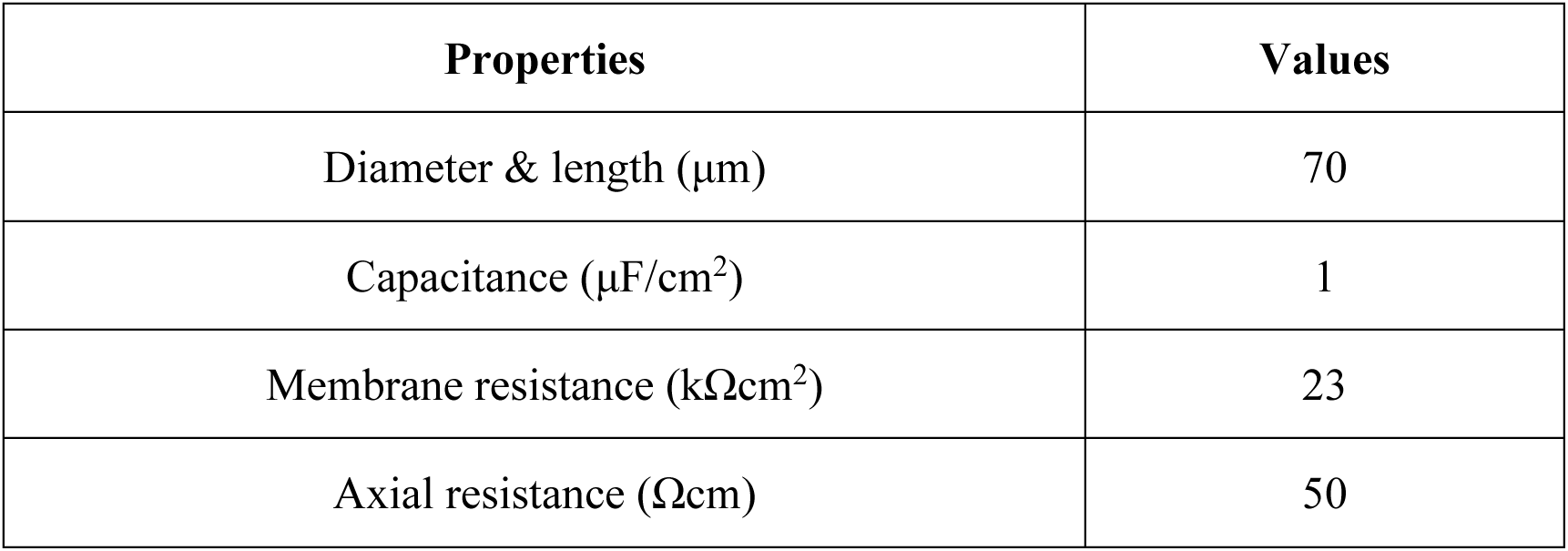
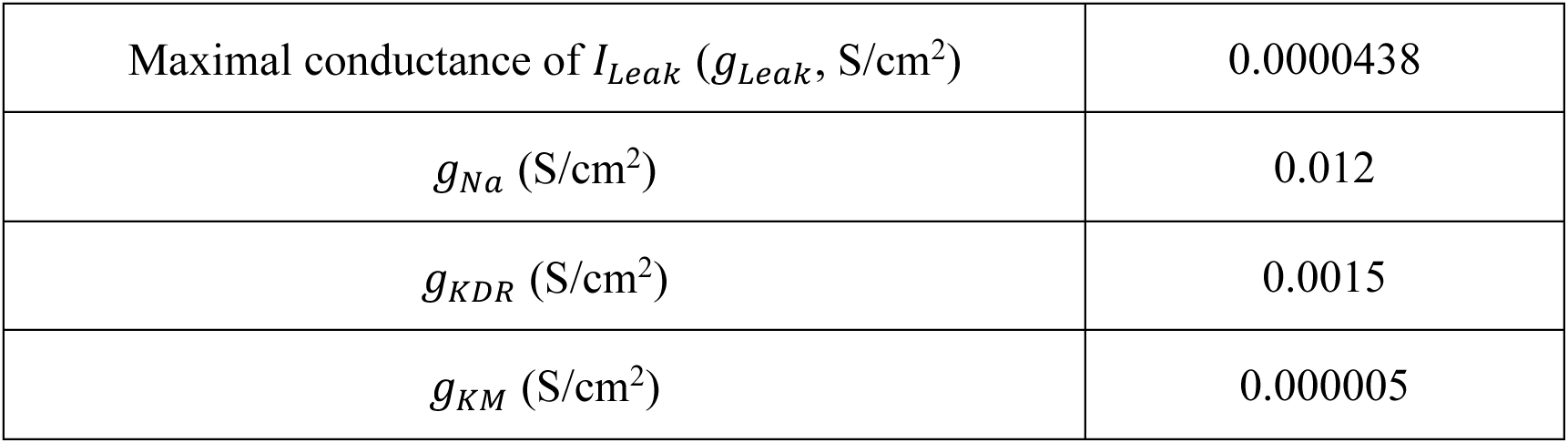
Membrane properties of the *in silico* boundary vector cell model.

| Properties | Values |
| --- | --- |
| Diameter & length ($\mu\text{m}$) | 70 |
| Capacitance ($\mu\text{F}/\text{cm}^2$) | 1 |
| Membrane resistance ($k\Omega\text{cm}^2$) | 23 |
| Axial resistance ($\Omega\text{cm}$) | 50 |
| Maximal conductance of $I_{Leak}$ ($g_{Leak}$, S/cm<sup>2</sup>) | 0.0000438 |
| $g_{Na}$ (S/cm<sup>2</sup>) | 0.012 |
| $g_{KDR}$ (S/cm<sup>2</sup>) | 0.0015 |
| $g_{KM}$ (S/cm<sup>2</sup>) | 0.000005 |

Boundary vector cell’s spike was evoked by giving a current input that is modulated by a Gaussian tuning curve^24,25^ using the following equation:

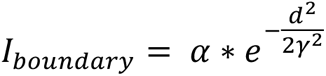

Where *I_boundary_* is the amount of input current to the soma of boundary vector cell model, *α* is a constant input current value (0.0525 nA), *d* is the distance from current position to the boundary in the environment, and *γ* is the boundary vector cell’s spatial receptive field size (10 cm). The peak spike firing rate of boundary vector cell model was set to 15 Hz, similar to *in vivo* observations^26^. To implement *in vivo*-like noise, boundary vector cell received Gaussian noise current input (mean = 0 nA, sd = 0.101 nA).

As RSC neuron has been reported to receive direct excitatory synaptic input from boundary vector cells^27^, RSC neuron was modeled to receive boundary vector cell spike-evoked excitatory postsynaptic current (*I_syn_*) using double-exponential conductance model^28^:

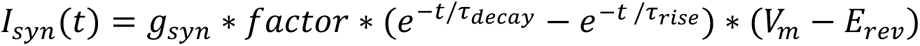

where *g_syn_* is the maximal conductance of synapse (0.001087 μS), *factor* is the normalizing constant, *τ_decay_* is the decay time constant (5 ms), *τ_rise_* is the rise time constant (4 ms), [*V_m_* is the membrane potential, and *E_rev_* is the reversal potential of the synapse model (0 mV). To simulate a simple vertex cell model receiving boundary vector cell inputs, we assumed that an *in silico* RSC neuron model received inputs from boundary vector cells that represent each edge of the open chamber. Thus, three, four, and six boundary vector cell-evoked subthreshold synaptic inputs were given to the RSC neuron model in triangle, square, and hexagon open chambers, respectively (Extended Data Fig. 16). All computational models were simulated using NEURON 8.2^29^ with a temperature of 30℃ and an integration time step of 0.1 ms. *In vivo* mouse trajectory data with linear interpolation (0.1 ms interval) was used to build 2D virtual environments for *in silico* simulation. Simulated spikes were analyzed using the same method that was used for analyzing *in vivo* experimental data.

#### Statistics

Statistical significance was measured using paired or unpaired Student’s *t*-test, or one-way ANOVA with *post hoc* Tukey’s test. For circular data, Wilcoxon signed-rank test^30,31^ were used. *p* value less than 0.05 was considered statistically significant.

---

**Extended Data Fig. 1.**
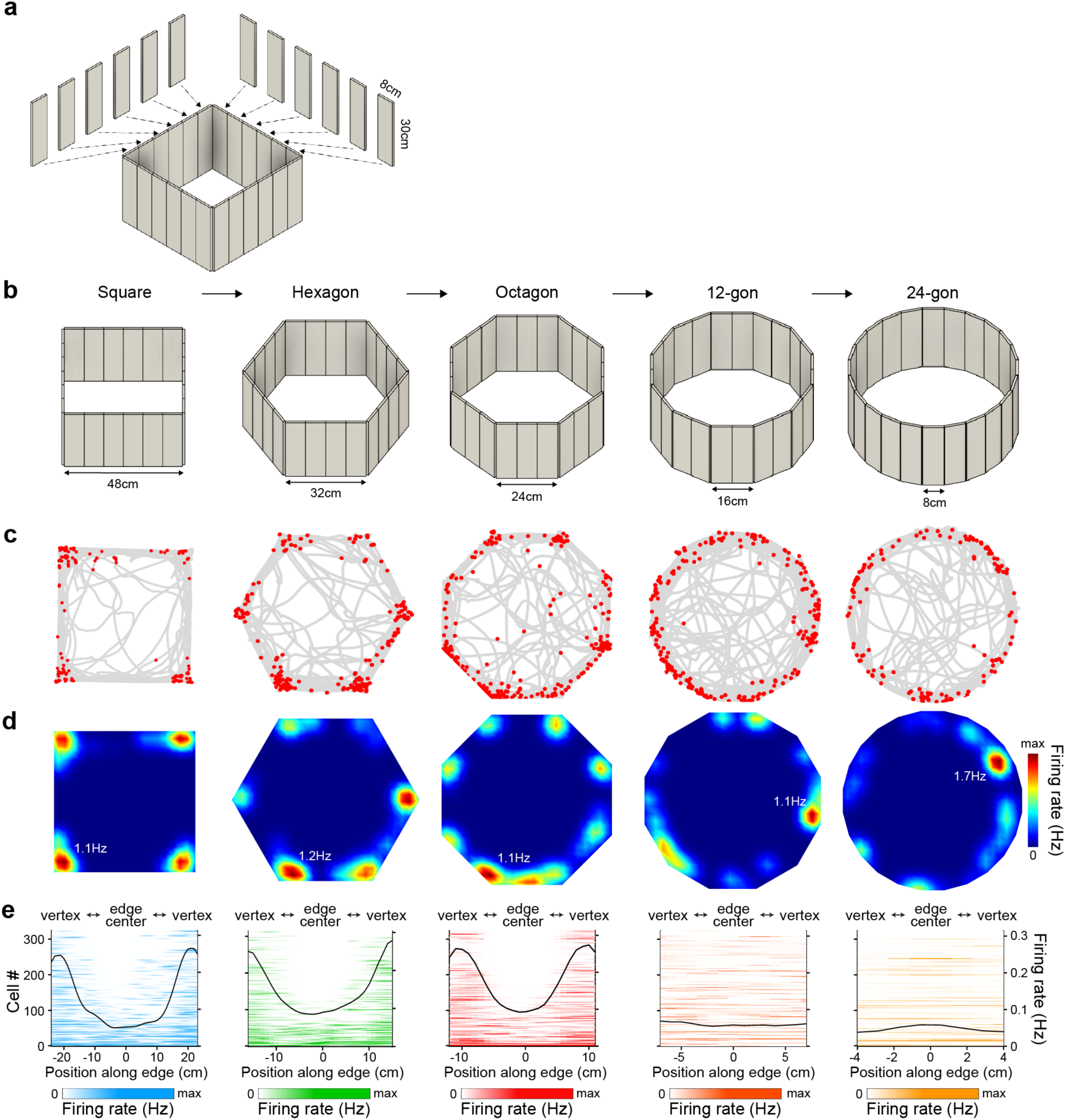
Vertex-selective neurons in the retrosplenial cortex in different shapes of morph open chambers. (**a**) Schematic illustration of constructing different shapes of regular polygon-shaped morph open chambers. (**b**) Morph open chambers were configured as square, hexagon, octagon, 12-gon, and 24-gon that had equal perimeter. (**c**, **d**) Representative spike-trajectory plot (c) showing the mouse trajectory (gray line) superimposed with spike locations (red dots) and the corresponding firing rate maps (d) of a single neuron in the granular retrosplenial cortex (RSC) during free exploration in each open chamber. Firing rate is color-coded (inset: maximum firing rate). (**e**) RSC neuronal firing rate map along the position on a given edge in each chamber. Black line: mean firing rate of all neurons (323 RSC neurons in 4 mice).

**Extended Data Fig. 2.**
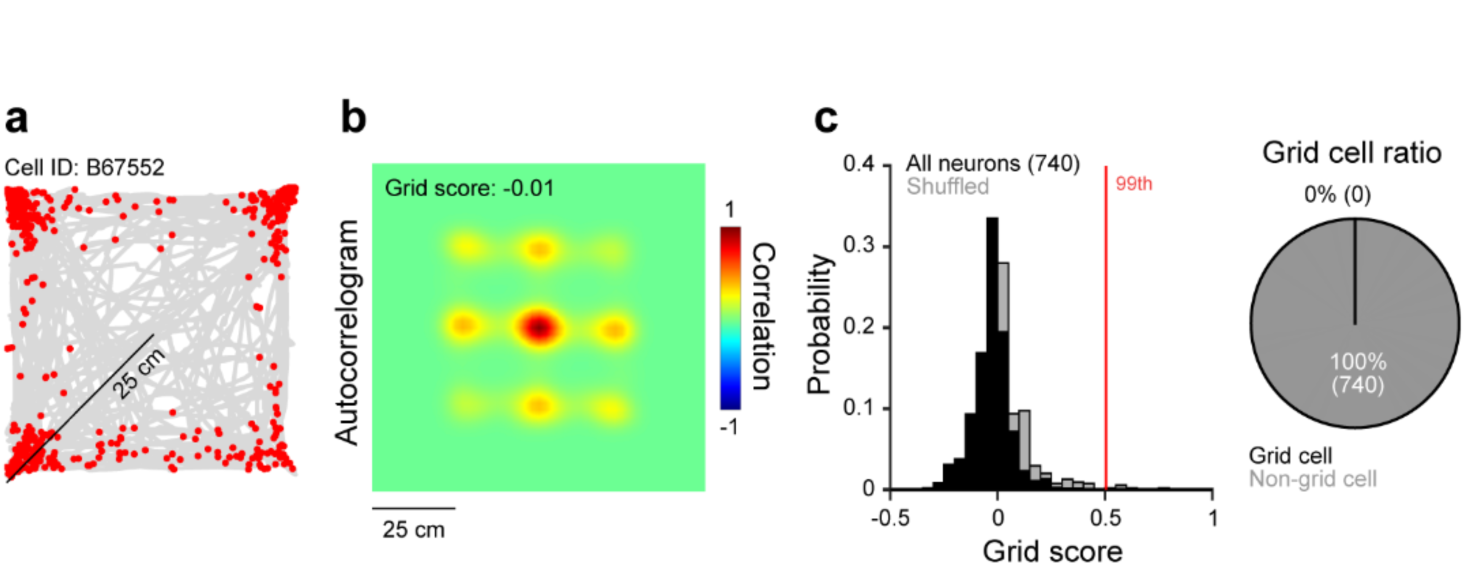
Grid scores of neurons in the retrosplenial cortex. (**a**, **b**) Representative spike-trajectory plot (a) showing the mouse trajectory (gray line) superimposed with spike locations (red dots) and the corresponding spatial autocorrelogram (b) of a single neuron in the granular retrosplenial cortex during free exploration in a square open chamber. Autocorrelation in (b) is color-coded (inset: grid score). (**c**) Probability distributions (left) of grid scores in a square open chamber (black) plotted together with the randomly shuffled grid scores for comparison (gray). Red line: 99^th^ percentile of randomly shuffled grid score distribution. Pie chart (right) showing ratio of grid cells and non-grid cells.

**Extended Data Fig. 3.**
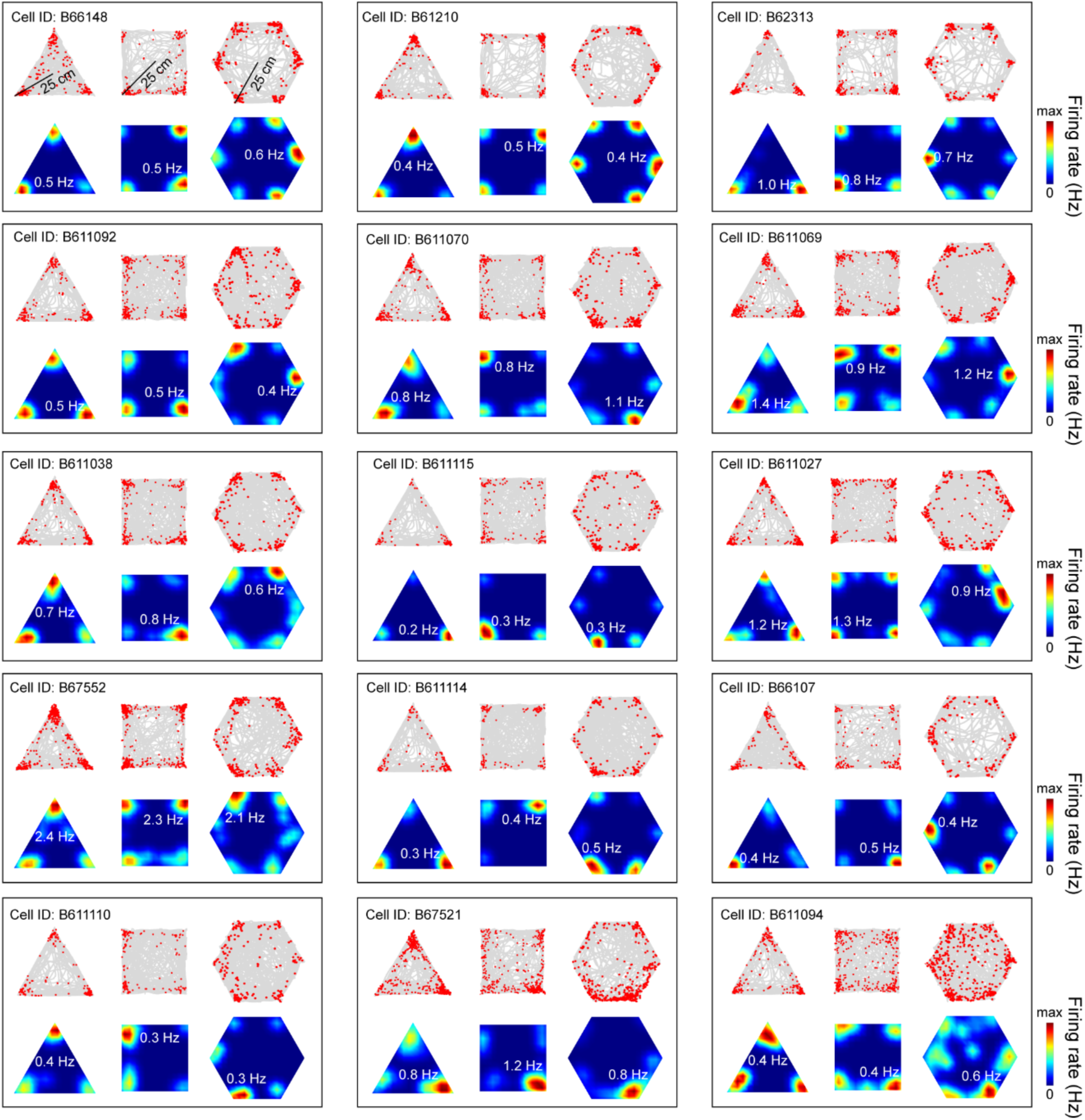
Examples of vertex cells in the retrosplenial cortex. Representative spike-trajectory plot (top) showing the mouse trajectory (gray line) superimposed with spike locations (red dots) and the corresponding firing rate maps (bottom) of vertex cells during free exploration in triangle, square, and hexagon open chambers. Examples of fifteen vertex cells in the granular retrosplenial cortex that sustained vertex-selective responses in all three open chambers are shown. Firing rate is color-coded (inset: maximum firing rate).

**Extended Data Fig. 4.**
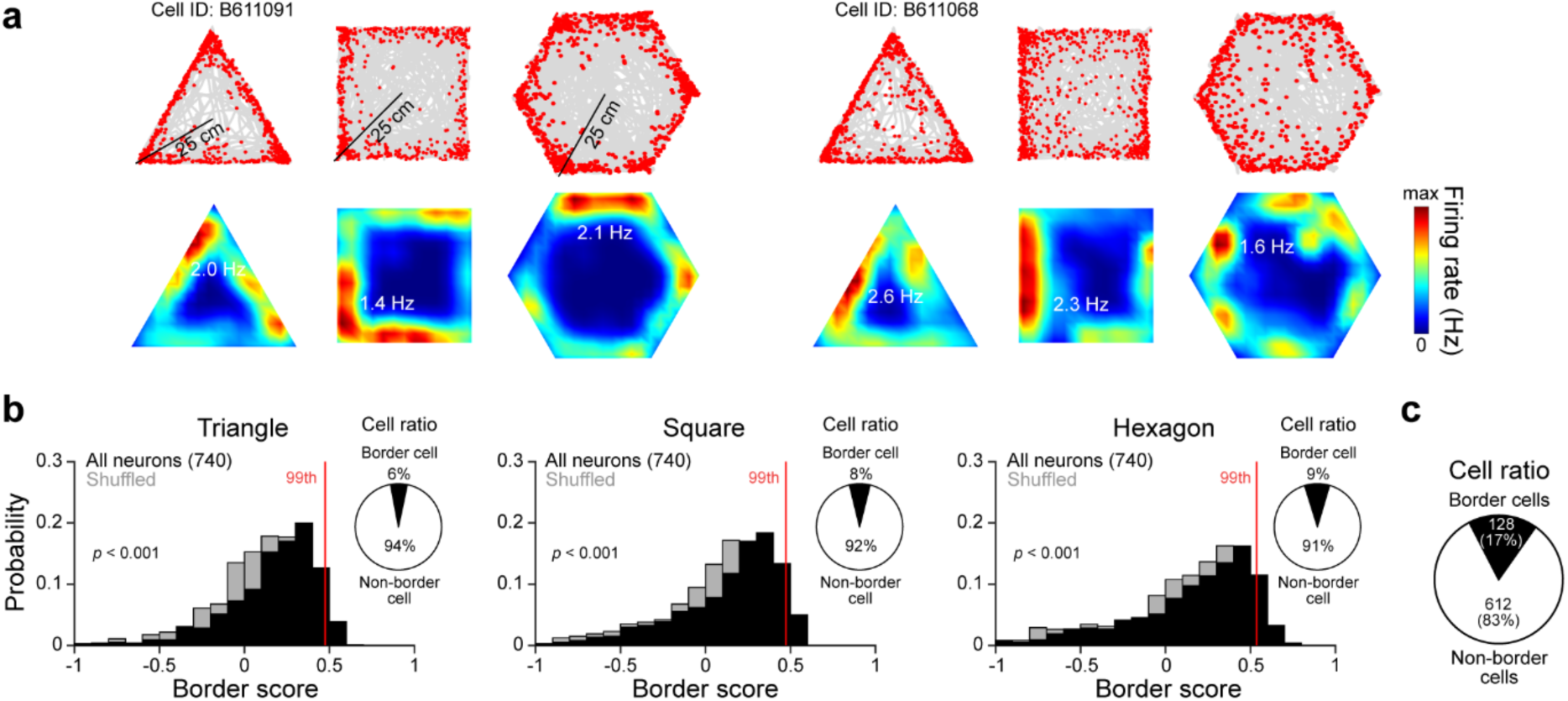
Border scores of neurons in the retrosplenial cortex. (**a**) Representative spike-trajectory plot (top) showing the mouse trajectory (gray line) superimposed with spike locations (red dots) and the corresponding firing rate maps (bottom) of two border cells in the granular retrosplenial cortex during free exploration in triangle, square, and hexagon open chambers. Firing rate is color-coded (inset: maximum firing rate). (**b**) Probability distributions of border scores in each open chamber (black) plotted together with the randomly shuffled border scores for comparison (gray). Red line: 99^th^ percentile of randomly shuffled border score distribution. Inset: pie chart showing ratio of border cells and non-border cells in each open chamber. (**c**) Pie chart showing border cell ratio across three open chambers. Unpaired Student’s *t* test (b).

**Extended Data Fig. 5.**
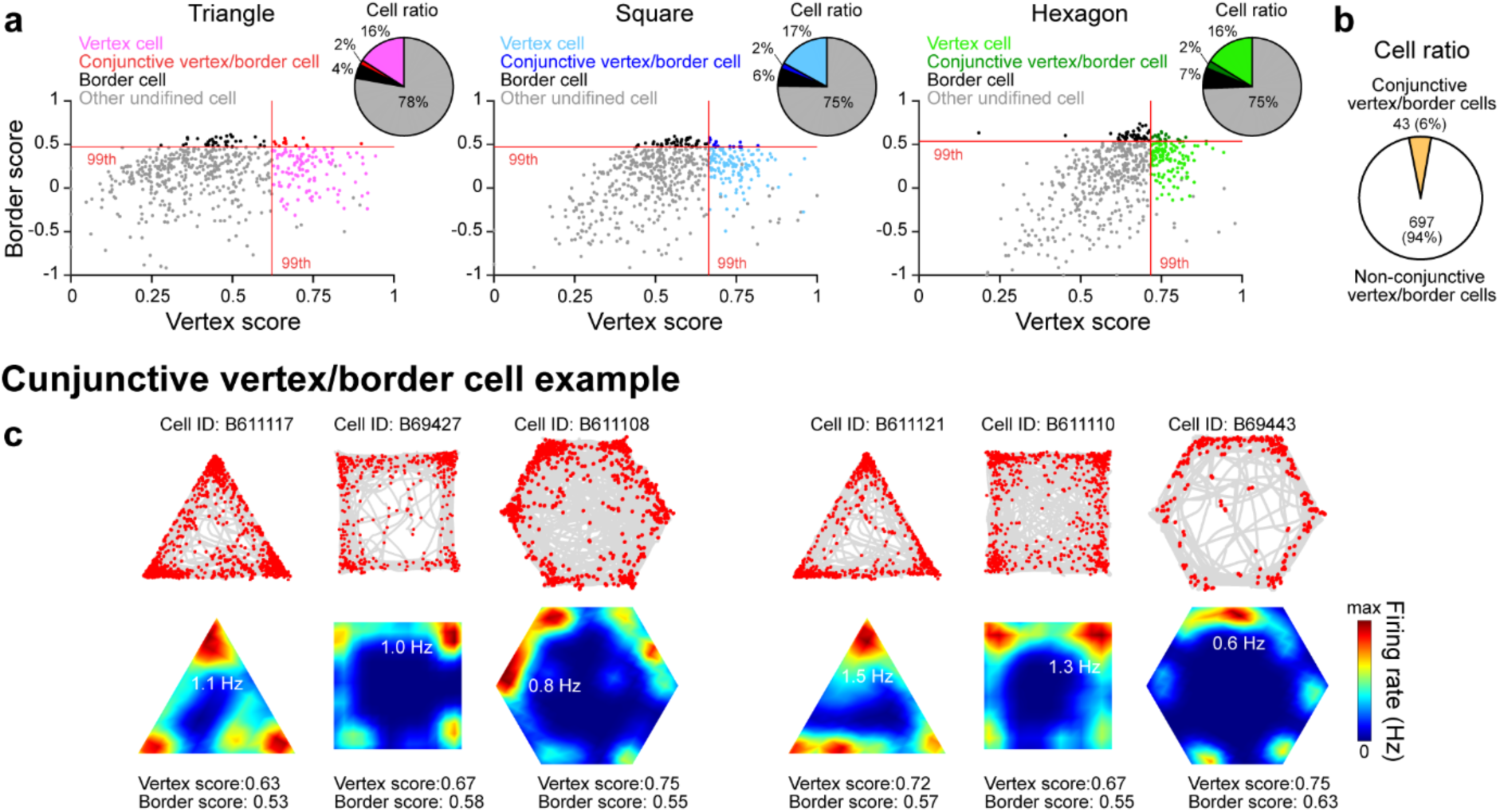
Conjunctive vertex/border cells in the retrosplenial cortex. (**a**) Scatter plot of border scores plotted as a function of vertex scores, which was used to classify vertex cells, conjunctive vertex/border cells, border cells, and other undefined cells. Red line: 99^th^ percentile of randomly shuffled vertex score (vertical) and border score (horizontal) distributions. Inset: pie chart showing ratio of vertex cells, conjunctive vertex/border cells, border cells, and other undefined cells in each open chamber. (**b**) Pie chart showing conjunctive vertex/border cell ratio across three open chambers. (**c**) Representative spike-trajectory plot (top) showing the mouse trajectory (gray line) superimposed with spike locations (red dots) and the corresponding firing rate maps (bottom) of two conjunctive vertex/border cells in the granular retrosplenial cortex during free exploration in triangle, square, and hexagon open chambers. Firing rate is color-coded (inset: maximum firing rate).

**Extended Data Fig. 6.**
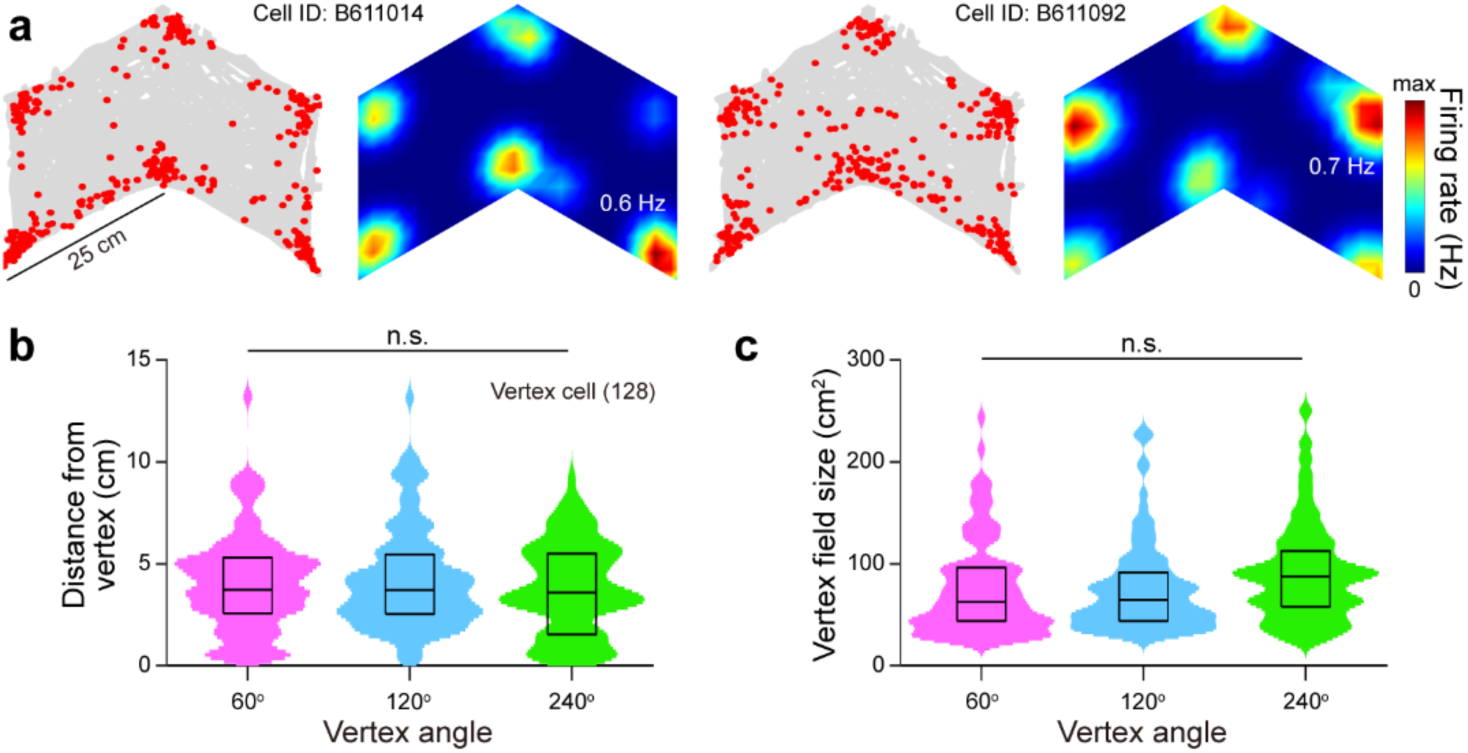
Vertex cell responses to a regular concave open chamber. (**a**) Representative spike-trajectory plot showing the mouse trajectory (gray line) superimposed with spike locations (red dots) and the corresponding firing rate maps of two vertex cells in the granular retrosplenial cortex during free exploration in a concave open chamber. Firing rate is color-coded (inset: maximum firing rate). (**b, c**) Violin plot showing distance of vertex fields from vertex (b) and the vertex field size (c) of vertex cells at different vertex angles (128 vertex cells in 8 mice). Box plot shows 25^th^, 50^th^, and 75^th^ percentile values. n.s.: *p* > 0.05 for one-way ANOVA with *post hoc* Tukey’s test (b, c).

**Extended Data Fig. 7.**
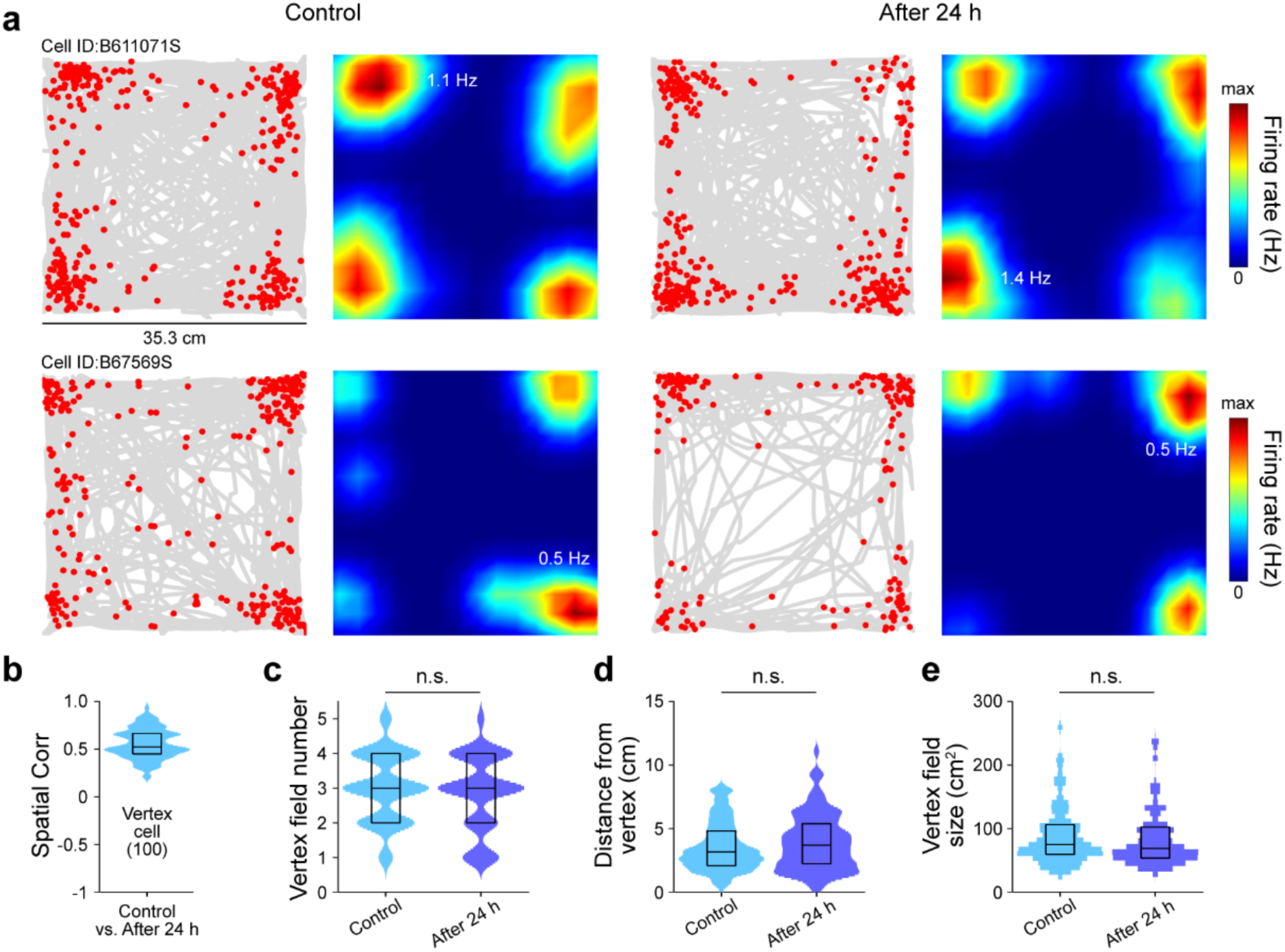
Vertex cell responses over time. (**a**) Representative spike-trajectory plot showing the mouse trajectory (gray line) superimposed with spike locations (red dots) and the corresponding firing rate maps of two vertex cells in the granular retrosplenial cortex during free exploration in a square open chamber (Control) and exploration of the same chamber after 24 h (After 24 h). Firing rate is color-coded (inset: maximum firing rate). (**b**) Violin plot showing spatial correlation coefficient (Spatial Corr) of vertex cells between the two conditions. (**c-e**) Violin plot showing the number of vertex fields (c), distance of vertex fields from vertex (d), and the vertex field size (e) of vertex cells in each condition (100 vertex cells in 7 mice). Box plot shows 25^th^, 50^th^, and 75^th^ percentile values. n.s.: *p* > 0.05 for paired Student’s *t* test (c-e).

**Extended Data Fig. 8.**
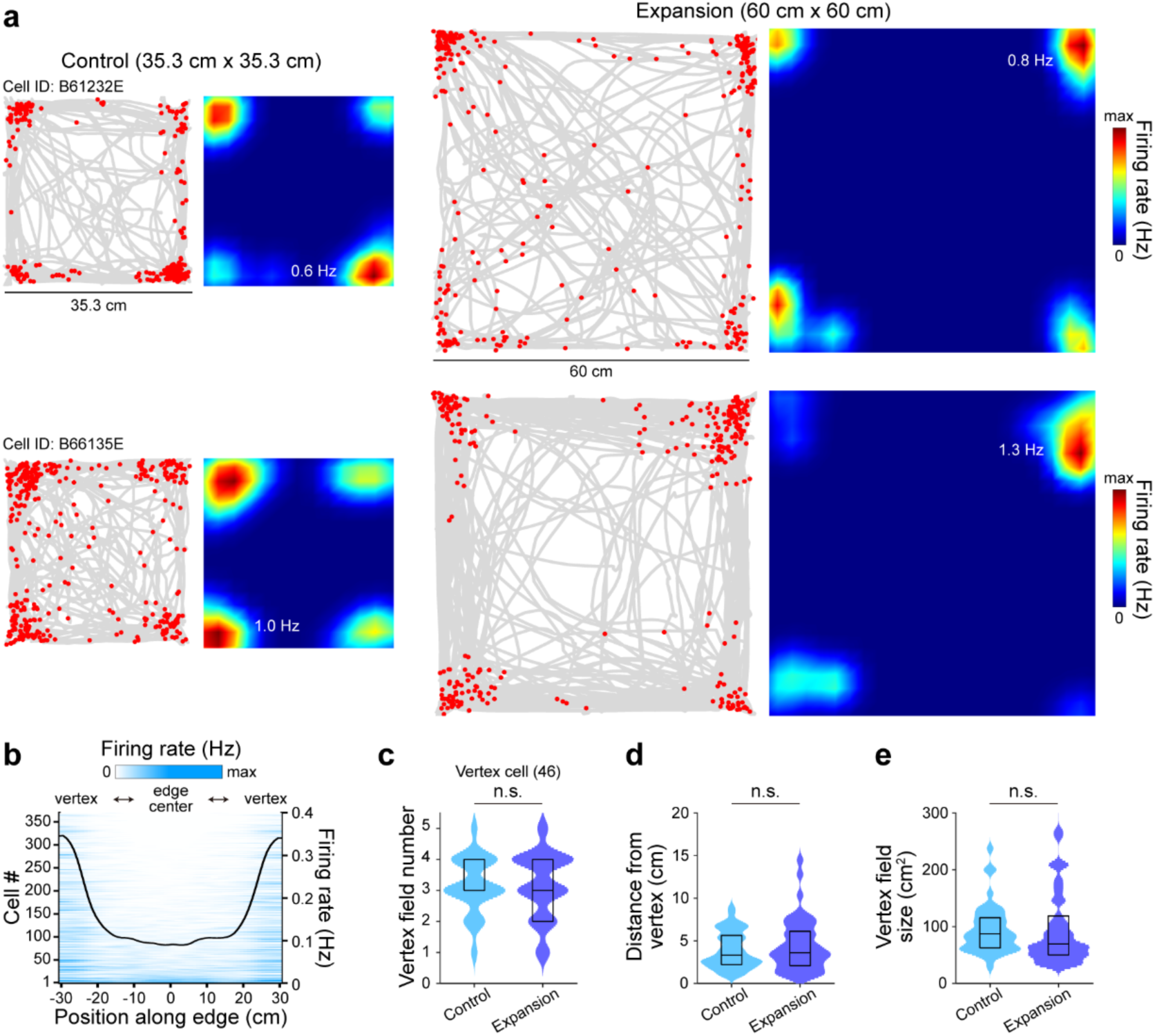
Vertex cell responses to chamber size expansion. (**a**) Representative spike-trajectory plot showing the mouse trajectory (gray line) superimposed with spike locations (red dots) and the corresponding firing rate maps of two vertex cells in the granular retrosplenial cortex (RSC) during free exploration in a square open chamber (Control, 35.3 cm x 35.3 cm) and an expanded square open chamber (Expansion, 60 cm x 60 cm). Firing rate is color-coded (inset: maximum firing rate). (**b**) RSC neuronal firing rate map along the position on a given edge in the expanded square open chamber. Black line: mean firing rate of all neurons. (**c-e**) Violin plot showing the number of vertex fields (c), distance of vertex fields from vertex (d), and the vertex field size (e) of vertex cells in each condition (46 vertex cells in 8 mice). Box plot shows 25^th^, 50^th^, and 75^th^ percentile values. n.s.: *p* > 0.05 for paired Student’s *t* test (c-e).

**Extended Data Fig. 9.**
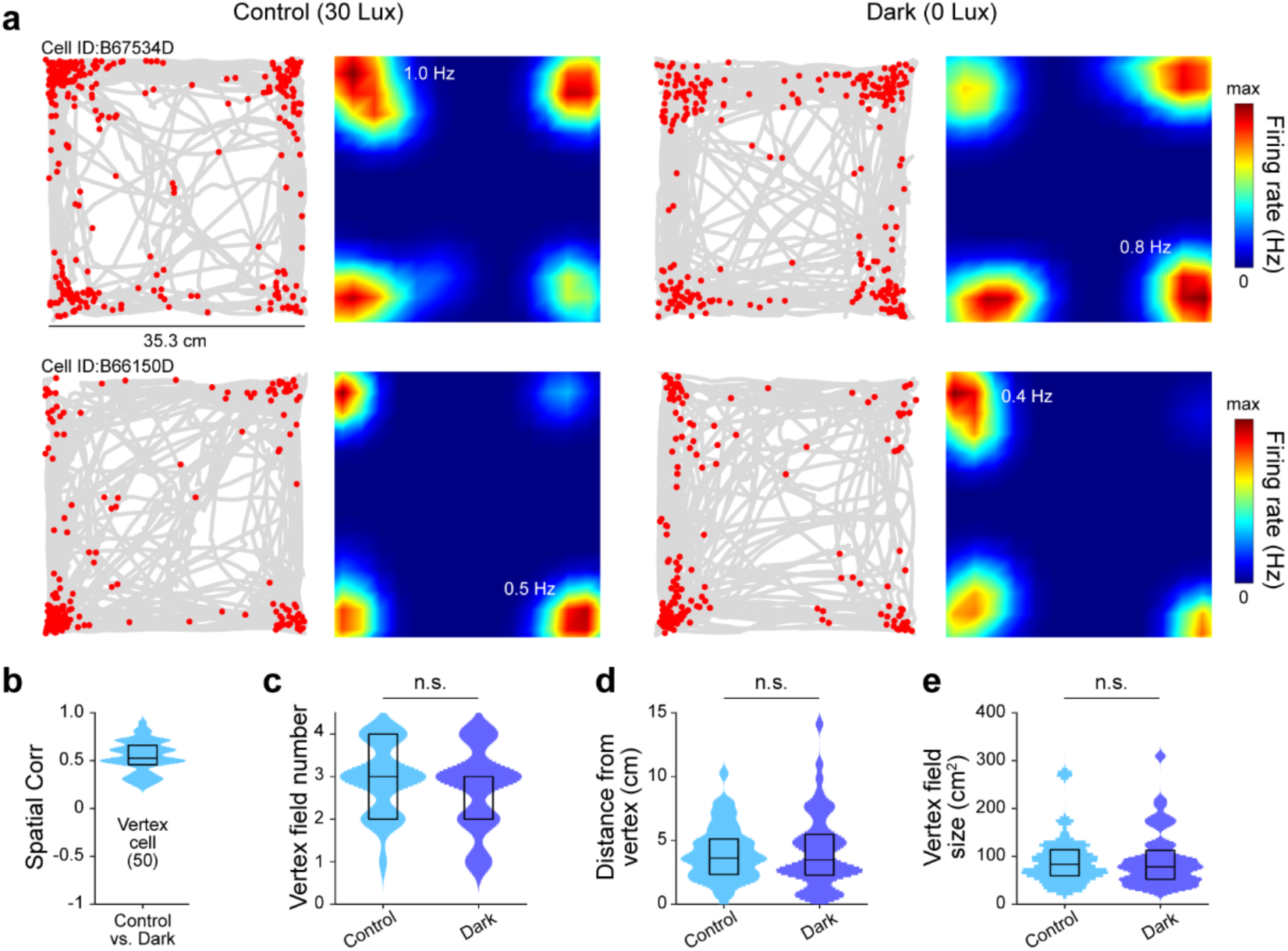
Vertex cell responses in dark condition. (**a**) Representative spike-trajectory plot showing the mouse trajectory (gray line) superimposed with spike locations (red dots) and the corresponding firing rate maps of two vertex cells in the granular retrosplenial cortex during free exploration in a square open chamber with (Control, 30 Lux) and without visible light (Dark, 0 Lux). Firing rate is color-coded (inset: maximum firing rate). (**b**) Violin plot showing spatial correlation coefficient (Spatial Corr) of vertex cells between the two conditions. (**c-e**) Violin plot showing the number of vertex fields (c), distance of vertex fields from vertex (d), and the vertex field size (e) of vertex cells in each condition (50 vertex cells in 4 mice). Box plot shows 25^th^, 50^th^, and 75^th^ percentile values. n.s.: *p* > 0.05 for paired Student’s *t* test (c-e).

**Extended Data Fig. 10.**
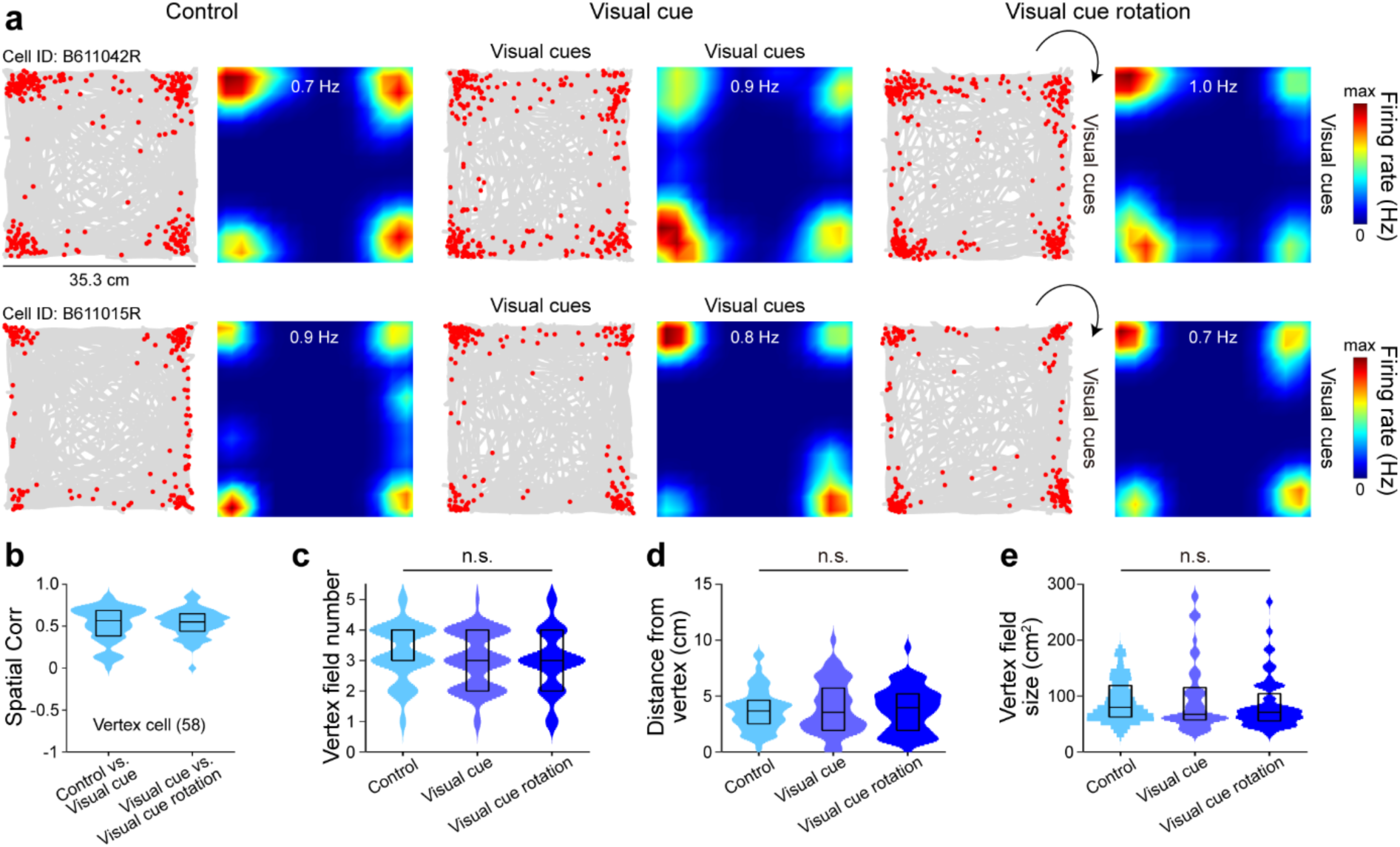
Vertex cell responses to visual cue rotation. (**a**) Representative spike-trajectory plot showing the mouse trajectory (gray line) superimposed with spike locations (red dots) and the corresponding firing rate maps of two vertex cells in the granular retrosplenial cortex during free exploration in a square open chamber without (Control), with visual cues introduced (Visual cue), and with 90° rotated visual cue on other side of the chamber (Visual cue rotation). Firing rate is color-coded (inset: maximum firing rate). (**b**) Violin plot showing spatial correlation coefficient (Spatial Corr) of vertex cells between the three conditions. (**c-e**) Violin plot showing the number of vertex fields (c), distance of vertex fields from vertex (d), and the vertex field size (e) of vertex cells in each condition (58 vertex cells in 6 mice). Box plot shows 25^th^, 50^th^, and 75^th^ percentile values. n.s.: *p* > 0.05 for one-way ANOVA with *post hoc* Tukey’s test (c-e).

**Extended Data Fig. 11.**
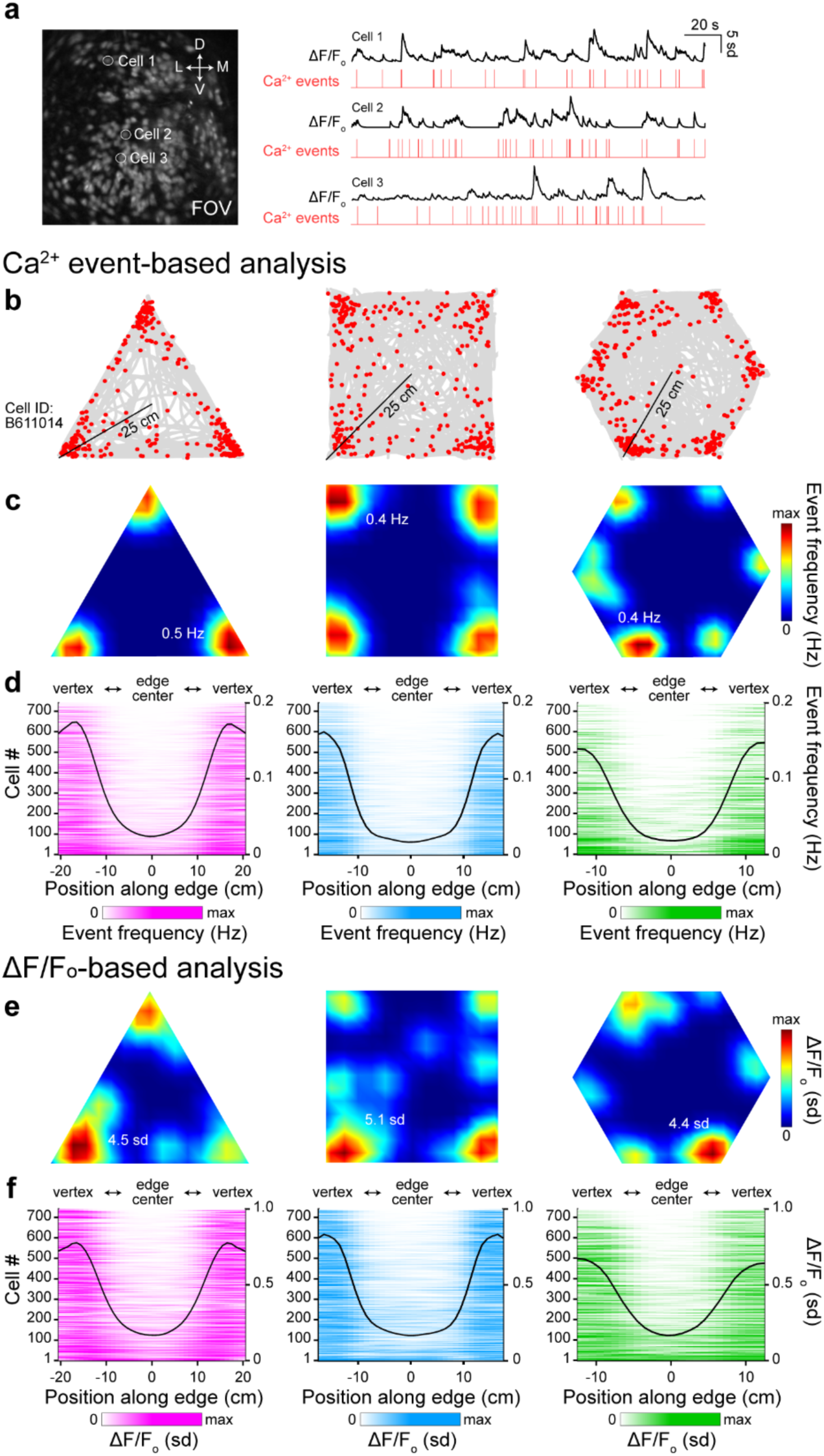
Vertex-selective neuronal responses in the retrosplenial cortex captured by Ca^2+^ event- or ΔF/Fo-based analysis. (**a**) Example field of view (FOV) of GCaMP6s-expressing neurons in the granular retrosplenial cortex (RSC) during Ca^2+^ imaging (left). Representative raw Ca^2+^ signals (right, black) from which Ca^2+^ events were extracted (right, red). (**b**, **c**) Representative Ca^2+^ events-trajectory plot (b) showing the mouse trajectory (gray line) superimposed with Ca^2+^ event locations (red dots) and the corresponding firing rate maps (c) of the same RSC neuron in Fig. 1C, D during free exploration in triangle, square, and hexagon open chambers. Event frequency in (c) is color-coded (inset: maximum event frequency). (**d**) RSC neuronal event frequency map along the position on a given edge in each chamber. Black line: mean event frequency of all neurons. (**e, f**) Same with (c, d) but using raw ΔF/Fo signal.

**Extended Data Fig. 12.**
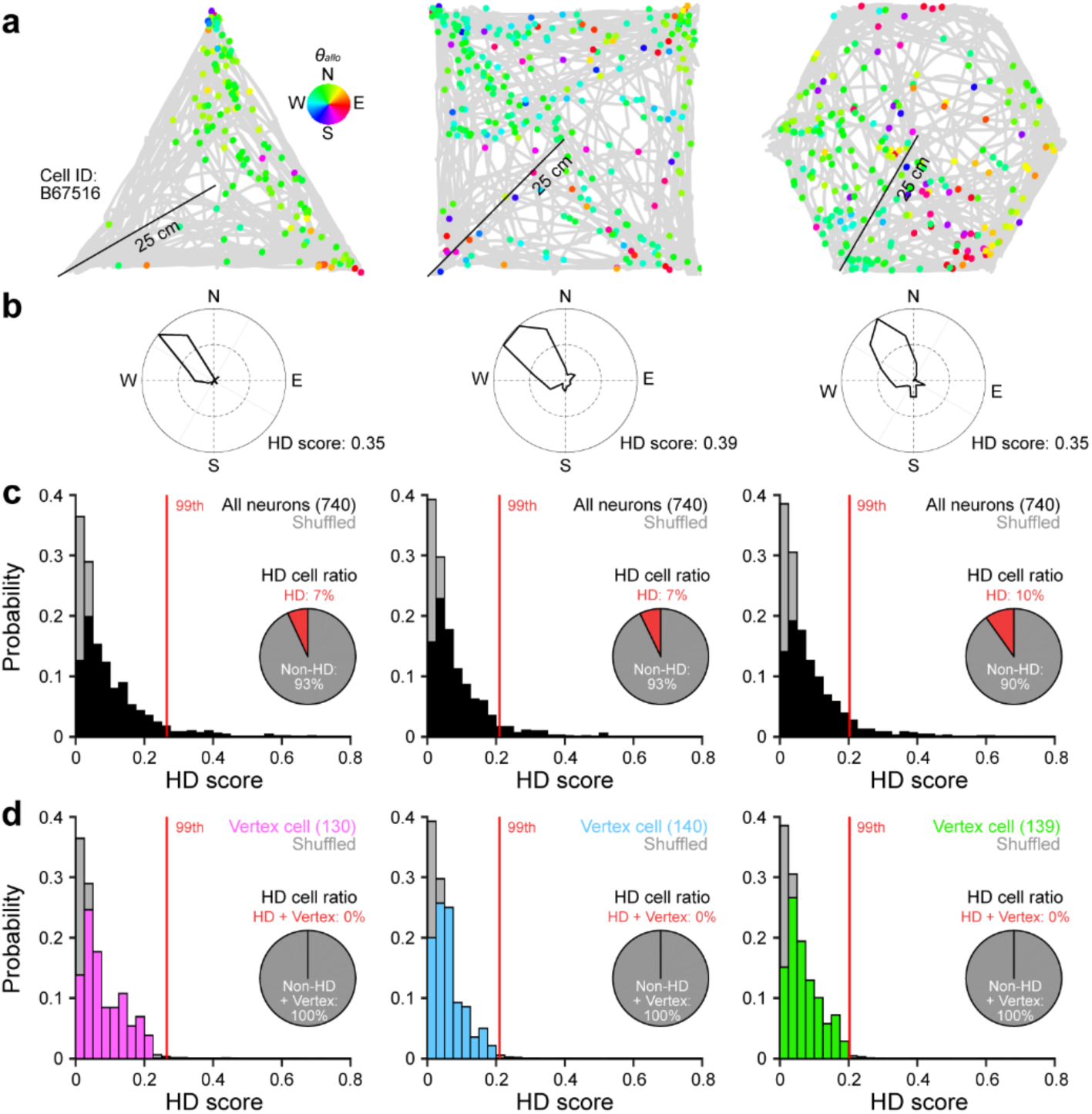
Head direction scores of neurons in the retrosplenial cortex. (**a**) Representative spike-trajectory plot showing the mouse trajectory (gray line) superimposed with spike locations (colored dots) of a single head direction cell (HD cell) in the granular retrosplenial cortex (RSC) during free exploration in triangle, square, and hexagon open chambers. Each spike is color-coded according to the corresponding allocentric head direction (*θallo*) (inset, North: N, East: E, South: S, and West: W). (**b**) The corresponding allocentric head direction tuning curves of the same HD cell in (a). Insets: head direction score (HD score). (**c**) Probability distributions of HD score of all RSC neurons in triangle, square, and hexagon open chambers (black) plotted together with the randomly shuffled HD scores for comparison (gray). Red line: 99^th^ percentile of randomly shuffled HD score distribution. Inset: pie chart showing ratio of HD cells and non-HD cells in each open chamber. (**d**) Same with (c) but for vertex cells.

**Extended Data Fig. 13.**
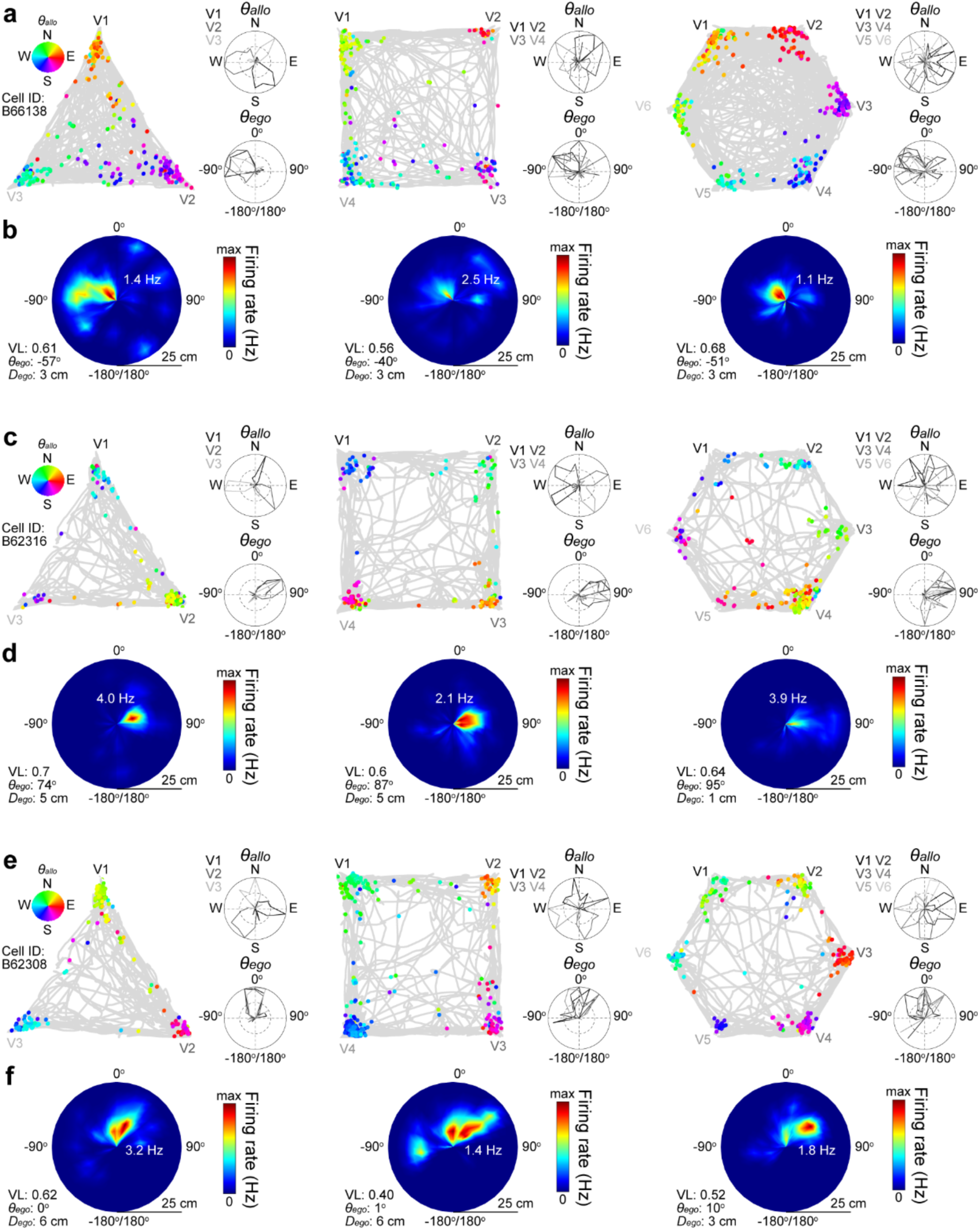
Examples of egocentric vertex vector cells in the retrosplenial cortex. (**a**) Representative spike-trajectory plot showing the mouse trajectory (gray line) superimposed with spike locations (colored dots) of a single egocentric vertex vector (EVV) cell in the granular retrosplenial cortex that sustained egocentric vector coding of vertices during free exploration in all triangle, square, and hexagon open chambers. Each spike is color-coded according to the corresponding allocentric head direction (*θallo*) (inset, North: N, East: E, South: S, and West: W). Allocentric head direction tuning curves and egocentric bearing tuning curves are color-coded for the corresponding closest vertex (Vi: *i*th vertex) in all open chambers. (**b**) The corresponding egocentric vertex rate map of the same EVV cell in (a). Firing rate is color-coded. Insets: mean resultant vector length (VL), preferred egocentric bearing (*θego*), preferred egocentric distance (*Dego*), and maximum firing rate. (**c-f**) Same with (a-b) but for other two EVV cells that sustained egocentric vector coding of vertices in all three open chambers.

**Extended Data Fig. 14.**
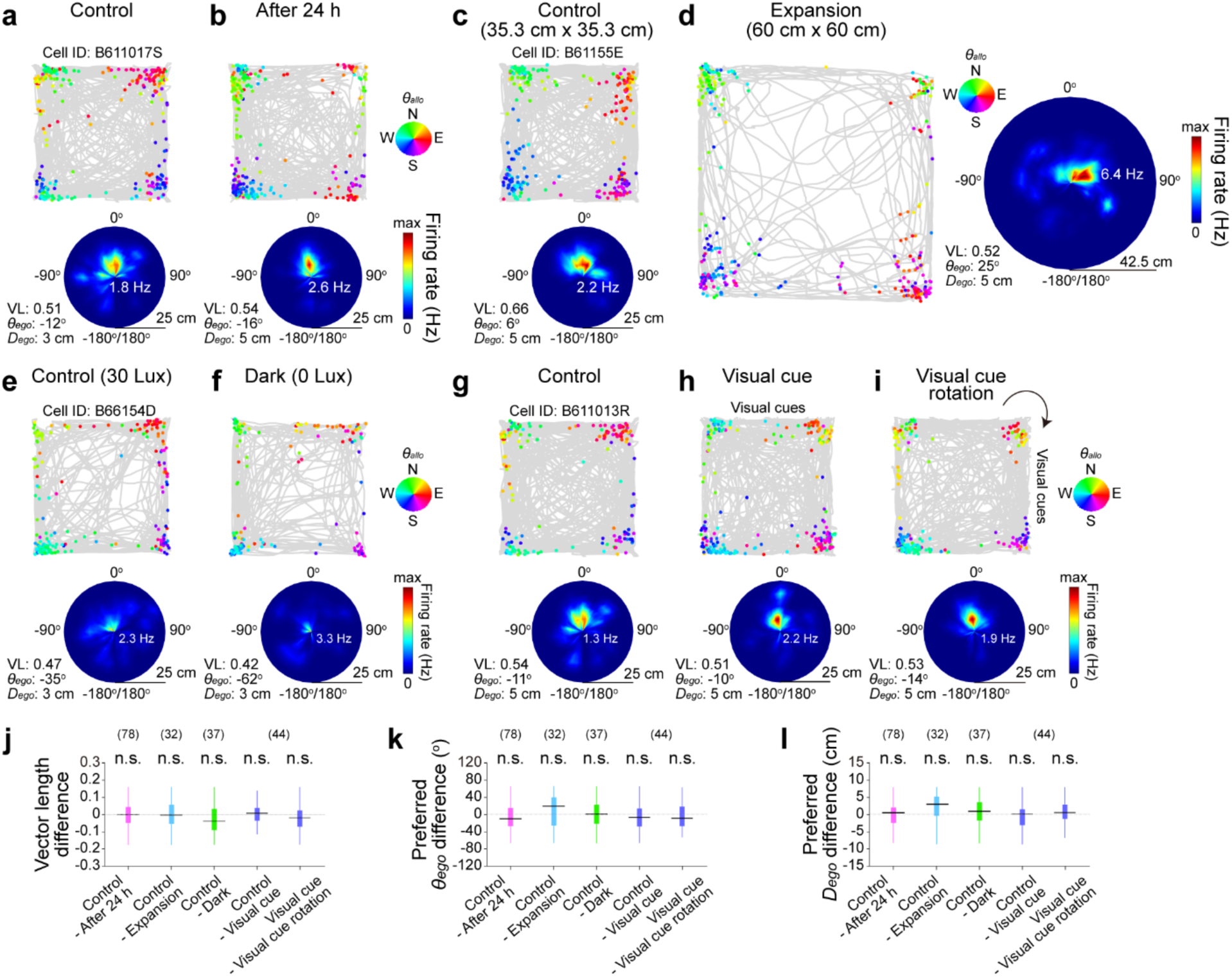
Stability of egocentric vector coding of vertex in the retrosplenial cortex across different conditions. (**a**-**i**) Representative spike-trajectory plot showing the mouse trajectory (gray line) superimposed with spike locations (colored dots) of egocentric vertex vector (EVV) cells in the granular retrosplenial cortex during free exploration in a square open chamber (a, c, e, g, Control) and exploration of the same chamber after 24 h (b, After 24 h), or an expanded square open chamber (d, Expansion), or the same chamber in the absence of visible light (f, Dark), or the same chamber with visual cue introduced (h, Visual cue), or with 90° rotated visual cue on other side of the chamber (i, Visual cue rotation). Each spike is color-coded according to the corresponding allocentric head direction (*θallo*) (inset, North: N, East: E, South: S, and West: W). The corresponding egocentric vertex rate map of the same EVV cells in spike-trajectory plot. Firing rate is color-coded. Insets: mean resultant vector length (VL), preferred egocentric bearing (*θego*), preferred egocentric distance (*Dego*), and maximum firing rate. (**j**-**l**) Box plot showing differences in VL (j), preferred *θego* (k), and *Dego* (l) of EVV cells across conditions. Box plot shows 25^th^, 50^th^, and 75^th^ percentile values. n.s.: *p* > 0.05 for paired Student’s *t* test (j, l). n.s.: *p* > 0.05 for Wilcoxon signed-rank test (k).

**Extended Data Fig. 15.**
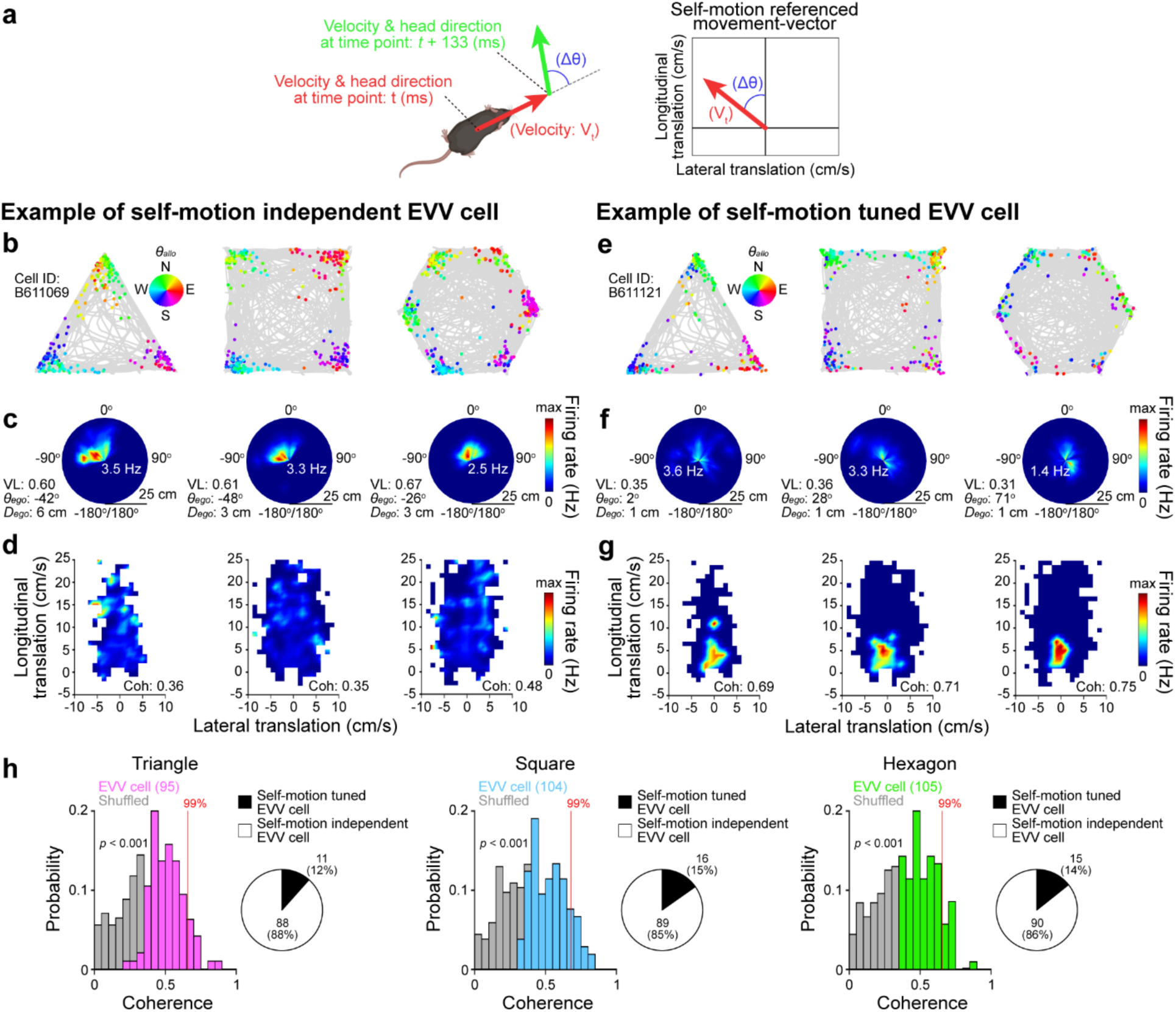
Self-motion tuning analysis of egocentric vertex vector cells in the retrosplenial cortex. (**a**) Schematic illustration of the mouse velocity and head direction at two example positions (left). Corresponding self-motion referenced movement-vector (right). (**b**) Representative spike-trajectory plot showing the mouse trajectory (gray line) superimposed with spike locations (colored dots) of a single egocentric vertex vector (EVV) cell in the granular retrosplenial cortex that is self-motion independent during free exploration in triangle, square, and hexagon open chambers. Each spike is color-coded according to the corresponding allocentric head direction (*θallo*) (inset, North: N, East: E, South: S, and West: W). (**c**) The corresponding egocentric vertex rate map of the same EVV cell in (b). Firing rate is color-coded. Insets: mean resultant vector length (VL), preferred egocentric bearing (*θego*), preferred egocentric distance (*Dego*), and maximum firing rate. (**d**) The corresponding self-motion referenced rate maps of the same EVV cell in (b) showing self-motion independence in all three open chambers. Inset: coherence value (Coh) of the self-motion referenced rate maps. (**e-g**) Same with (b-d) but of self-motion tuned EVV cell. (**h**) Probability distributions of coherence value of self-motion referenced rate maps in triangle (pink), square (blue), and hexagon open chambers (green) plotted together with the randomly shuffled coherence values for comparison (gray). Red line: 99^th^ percentile of randomly shuffled coherence value distribution. Inset: pie chart showing ratio of self-motion tuned EVV cells and self-motion independent EVV cells in each open chamber. Unpaired Student’s *t* test (h).

**Extended Data Fig. 16.**
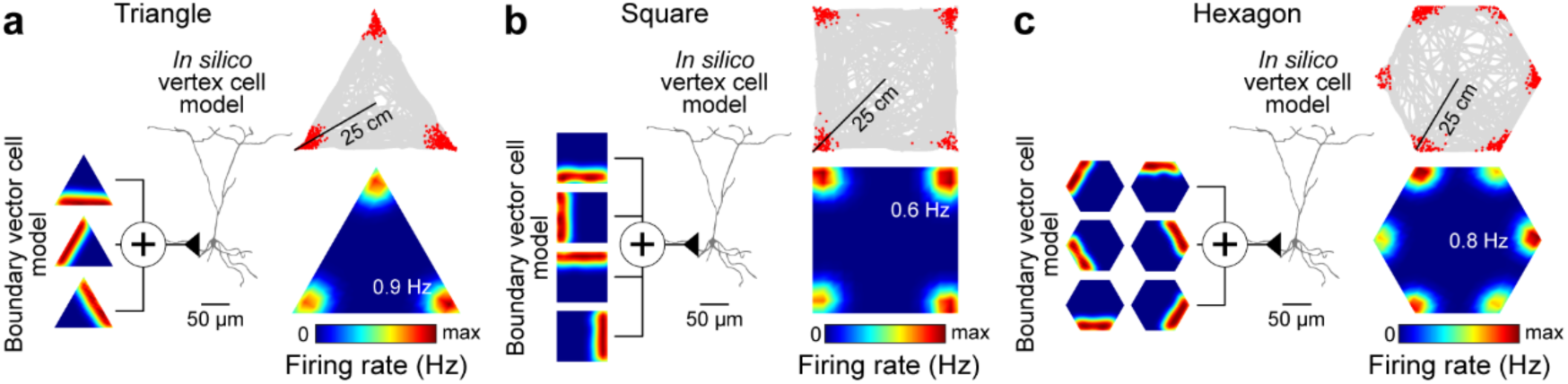
*In silico* modeling of vertex cells in the retrosplenial cortex. (**a**-**c**) Illustrations of a simple *in silico* computational model of vertex cell receiving boundary vector cell inputs in triangle (a), square (b), and hexagon open chambers (c).

**Extended Data Fig. 17.**
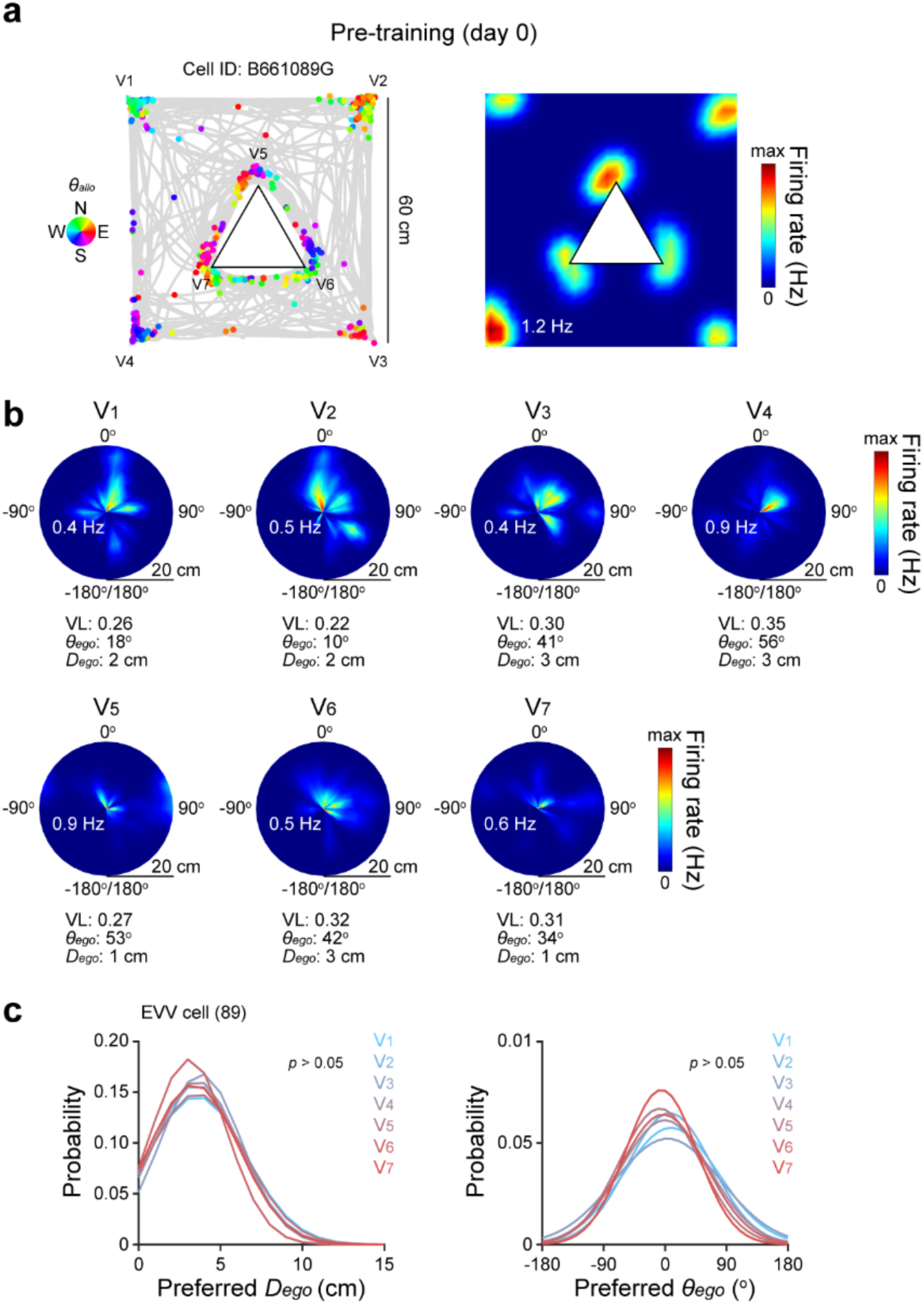
Robust egocentric vector coding of vertices of the open chamber and inner walls in the retrosplenial cortex. (**a**) Representative spike-trajectory plot (top) showing the mouse trajectory (gray line) superimposed with spike locations (colored dots) and the corresponding firing rate maps (bottom) of a single egocentric vertex vector (EVV) cell in the granular retrosplenial cortex during free exploration in a complex environment consisting of a square open chamber with a triangle-shaped inner walls (Vi: *i*th vertex in the environment). Each spike is color-coded according to the corresponding allocentric head direction (*θallo*) (inset, North: N, East: E, South: S, and West: W). Firing rate is color-coded (inset: maximum firing rate). (**b**) The corresponding egocentric vertex rate map (right) of the same EVV cell at each vertex of the open chamber and inner walls of. Firing rate is color-coded. Insets: mean resultant vector length (VL), preferred egocentric bearing (*θego*), preferred egocentric distance (*Dego*), and maximum firing rate. (**c**) Probability distribution of preferred *Dego* (left) and *θego* (right) of EVV cells at each vertex. Paired Student’s *t* test (c, left). Wilcoxon signed-rank test (c, right).

**Extended Data Fig. 18.**
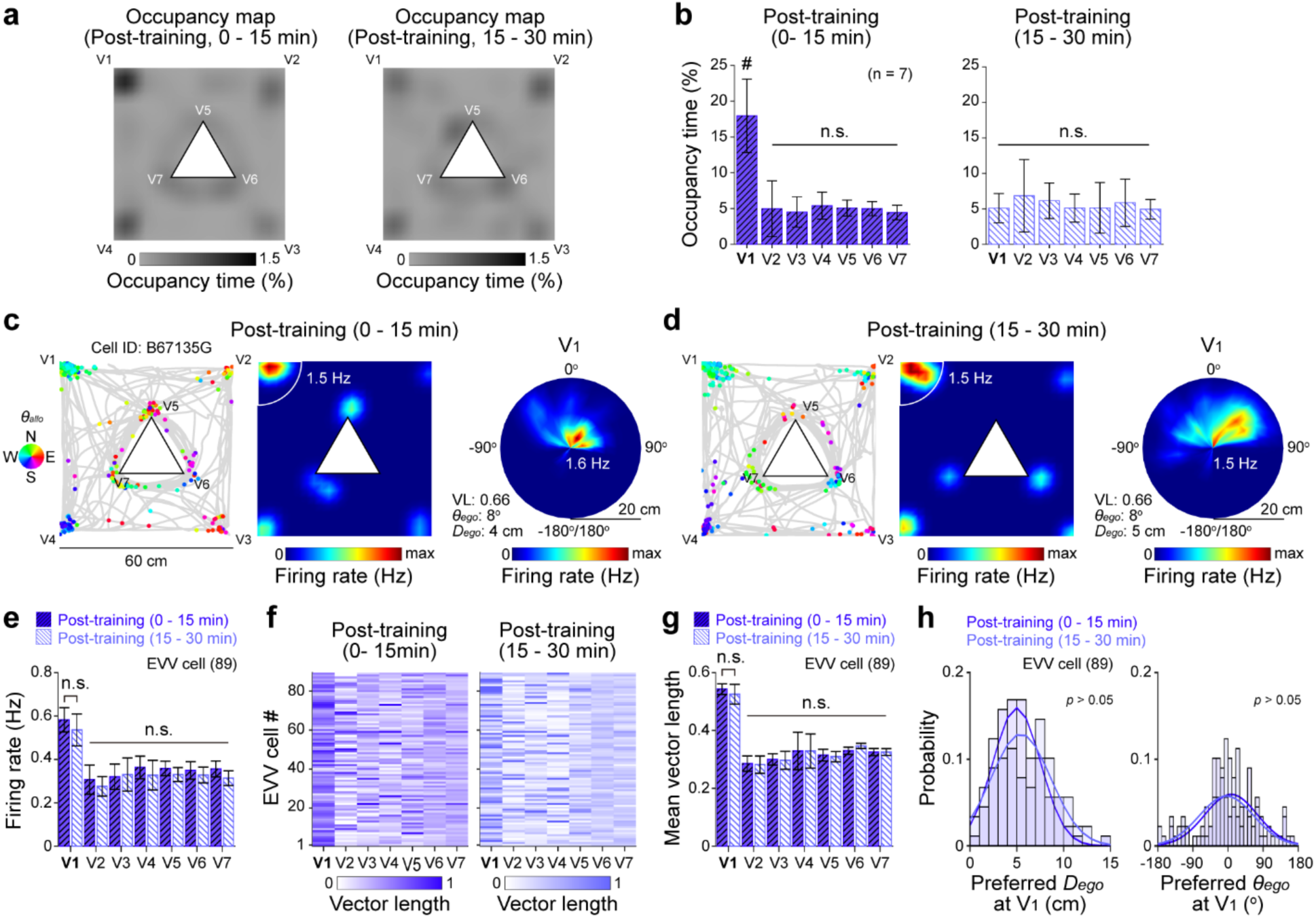
Occupancy time-independence of goal-directed navigation-induced strengthening of egocentric vector coding of geometric vertices in the retrosplenial cortex. (a) Representative occupancy map showing the mouse occupancy time distribution during free exploration in a square open chamber with a triangle-shaped inner walls (Vi: *i*th vertex) at the first half (left, 0 – 15 min) and second half (right, 15 – 30 min) of 30 min-long post-training session of goal-directed navigation task. Occupancy time is color-coded. (b) Mean occupancy time at each vertex (n = 7 mice). (c, d) Representative spike-trajectory plot (left) showing the mouse trajectory (gray line) superimposed with spike locations (colored dots) and the corresponding firing rate maps (middle) of the same egocentric vertex vector (EVV) cell in Fig. 4c, d during free exploration in the square open chamber with a triangle-shaped inner wall at the first half (c, 0 – 15 min) and the second half (d, 15 – 30 min) of post-training session. Each spike is color-coded according to the corresponding allocentric head direction (*θallo*) (inset, North: N, East: E, South: S, and West: W). Firing rate is color-coded (inset: maximum firing rate). The corresponding egocentric vertex rate map (right) of the same EVV cell at the vertex near goal location (V1, white line in firing rate map). Firing rate is color-coded. Insets: mean resultant vector length (VL), preferred egocentric bearing (*θego*), preferred egocentric distance (*Dego*), and maximum firing rate. (e-g) Average firing rate (e), VL map (f), and mean VL (g) of all EVV cells at each vertex in each condition (Post-training (0 – 15 min): blue, Post-training (15 – 30 min): dark blue). VL in (g) is color-coded. (h) Probability distribution of preferred *Dego* (left) and *θego* (right) of EVV cells at V1 in each condition. Bar graph shows mean ± SEM. #: *p* < 0.05, n.s.: *p* > 0.05 for one-way ANOVA with *post hoc* Tukey’s test (b). n.s.: *p* > 0.05 for paired Student’s *t* test (e, g, and h, left). Wilcoxon signed-rank test (h, right).

## Notes

### Competing Interest Statement

The authors have declared no competing interest.

